# Telomere-to-Telomere Phased Genome Assembly Using HERRO-Corrected Simplex Nanopore Reads

**DOI:** 10.1101/2024.05.18.594796

**Authors:** Dominik Stanojević, Dehui Lin, Sergey Nurk, Paola Florez de Sessions, Mile Šikić

**Affiliations:** Laboratory of AI in Genomics, Genome Institute of Singapore, A*STAR, Singapore, Singapore; Laboratory for Bioinformatics and Computational Biology, Faculty of Electrical Engineering and Computing, University of Zagreb, Zagreb, Croatia; Oxford Nanopore Technologies, Oxford, United Kingdom

**Author notes:** Corresponding Author: Mile Šikić.

## Abstract

Telomere-to-telomere phased assemblies have become the norm in genomics. To achieve these for diploid and even polyploid genomes, the contemporary approach involves a combination of two long-read sequencing technologies: high-accuracy long reads, e.g. Pacific Biosciences (PacBio) HiFi or Oxford Nanopore (ONT) ‘Duplex’ reads, and ultra-long ONT ‘Simplex’ reads. Using two different technologies increases the cost and the required amount of genomic DNA. Here, we show that comparable results are possible using error correction of ultra-long ONT Simplex reads and then assembling them using state-of-the-art de novo assembly methods. To achieve this, we have developed the deep learning-based HERRO framework, which corrects ONT Simplex reads while carefully preserving differences in related genomic sequences. Taking into account informative positions that differentiate the haplotypes or genomic repeat copies, HERRO achieves an increase of read accuracy of up to 100-fold for diploid human genomes. By combining HERRO with Verkko assembler, we achieve high contiguity on several human genomes by reconstructing many chromosomes telomere-to-telomere, including chromosomes X and Y. HERRO supports both R9.4.1 and R10.4.1 ONT Simplex reads and generalizes well to other species. These results provide an opportunity to reduce the cost of genome sequencing and use corrected ONT reads to analyze more complex genomes with different levels of ploidy or even aneuploidy.

## Introduction

Both accuracy and contiguity of genome assemblies have seen substantial improvements in recent years due to advancements in long-read sequencing technologies, new *de novo* assembly methods and dedicated work of the scientific community on manual curation. Resolving most chromosomes, haplotype-resolved telomere-to-telomere (T2T)^1^ diploid assemblies have become a standard.

To achieve haplotype-resolved T2T assemblies, it is currently necessary to use multiple technologies, primarily high-accuracy long reads such as HiFi reads produced by Pacific Biosciences sequencers or Duplex reads^2,3^ produced by Oxford Nanopore Technologies (ONT). ONT Simplex ultra-long (UL) reads, are also required. Moreover, chromosome-scale phasing usually requires either parental data^4^, single-cell DNA template strand sequencing (Strand-seq^5^) or cross-linking technology (such as Hi-C, Omni-C or Pore-C).

Using multiple technologies can significantly increase both the wet lab and dry lab workloads as well as the overall project cost. This approach requires a larger quantity of high molecular weight (HMW) gDNA, with separate library preparation and sequencing needed for each technology. Additionally, more complex algorithms are required to integrate reads from different technologies. Contemporary *de novo* assembly methods use high-accuracy long reads to construct initial assembly graphs, with lower-accuracy ONT Simplex reads longer than 100kb (known as ultra-long reads) serving as auxiliary data to simplify these graphs. Although PacBio HiFi and ONT Duplex reads offer high accuracy, they are usually insufficient for assembling complex regions, such as centromeres, on their own. They require the addition of ultra-long reads, which, even with their initial lower accuracy, are essential for successfully assembling these challenging regions and achieving haplotype-resolved T2T assemblies. Therefore, using ONT Simplex ultra-long reads as a single long-read sequencing technology can reduce the required gDNA, eliminate the need for additional DNA library preparation, streamline the sequencing workflow and unlock their full potential.

Several pipelines use error correction of noisy reads, such as ONT Simplex or PacBio CLR reads, at some stage to produce either collapsed, primary/alt or fully phased, haplotype-resolved assemblies. These include Canu^6^ + purge_dups^7^, FALCON-Unzip^8^, Flye^9^+HapDup^10,11^, Shasta^12^, and PECAT^13^. However, according to the results published in a recent study^13^, their performance is inferior compared to strategies that utilize both high-accuracy long reads and ultra-long ONT reads. For example, using recent ultra-long ONT data for the HG002 genome (Table 2), PECAT, the top performer among them, achieved a maximum (paternal/maternal) NG50 of 91.4/80.2 Mbp. In contrast, Hifiasm^14–16^ and Verkko^17^, utilizing multiple sequencing technologies – PacBio HiFi reads, ONT UL reads, and Illumina parental reads – produced assemblies with NG50 values exceeding 130 Mbp (ref. ^17^, Results). They also outperformed PECAT in phasing accuracy, as indicated by a considerably lower Hamming error rate (2.99%/1.49%^13^ per haplotype vs 0.5%^17^ total).

NextDenovo^18^, another recently published *de novo* assembler for noisy long reads, does not phase genomes. Hypo-assembler^19^ uses short Illumina reads to correct ONT reads.

Here, we present HERRO, a framework based on a deep learning model capable of correcting ONT Simplex reads. HERRO is optimized for both R9.4.1. and R10.4.1 pores and chemistry and supports both standard and ultra-long ONT Simplex reads.

HERRO achieves up to 100-fold improvement in read accuracy while preserving differences between related genomic sequences, e.g. heterozygous variants and differences between segmental duplications or satellite array copies. To accomplish this, we developed a novel deep learning model utilizing convolutional networks and self-attention architecture to process all-vs-all read overlap pile-ups. Model training has heavily benefitted from the availability of high-quality HG002 genome assembly (https://github.com/marbl/HG002).

Combining HERRO with existing de novo assemblers, such as Hifiasm, Verkko, and LJA^20^, we achieve highly contiguous haplotype-resolved assemblies, with many chromosomes represented as T2T contigs or scaffolds. Remarkably, despite being trained on only a few less complex chromosomes of the HG002 human genome, the HERRO model generalizes effectively to a wide range of organisms.

## Results

We developed HERRO, a method (Extended Data Fig. 1) and software package aimed at increasing the quality of ONT reads. HERRO is a read correction model that focuses on positions in reads representing differences between haplotypes or specific segments within segmental duplications. Since these positions hold valuable information for phasing, we refer to them as *informative positions*.

HERRO incorporates a newly developed deep learning model (Fig. 1) that combines convolutional blocks with a Transformer encoder. The model processes windows of stacked bases and quality scores extracted from all-vs-all pairwise alignments (obtained with minimap2^21^). Initially, the convolutional blocks process these windows to extract local information from each base and to aggregate information across different reads at the same position, generating representations for each read position. Subsequently, the model selectively processes *informative position candidates* – defined as those having at least two different bases (including gap symbol) that each appear a minimum of three times – through the Transformer encoder. This encoder is designed to learn interactions between these positions. By learning to predict bases at these challenging positions, HERRO accurately corrects reads while preserving the differences between haplotypes. For the remaining read positions, we use a simple majority voting method to predict the correct bases.

**Fig. 1.**
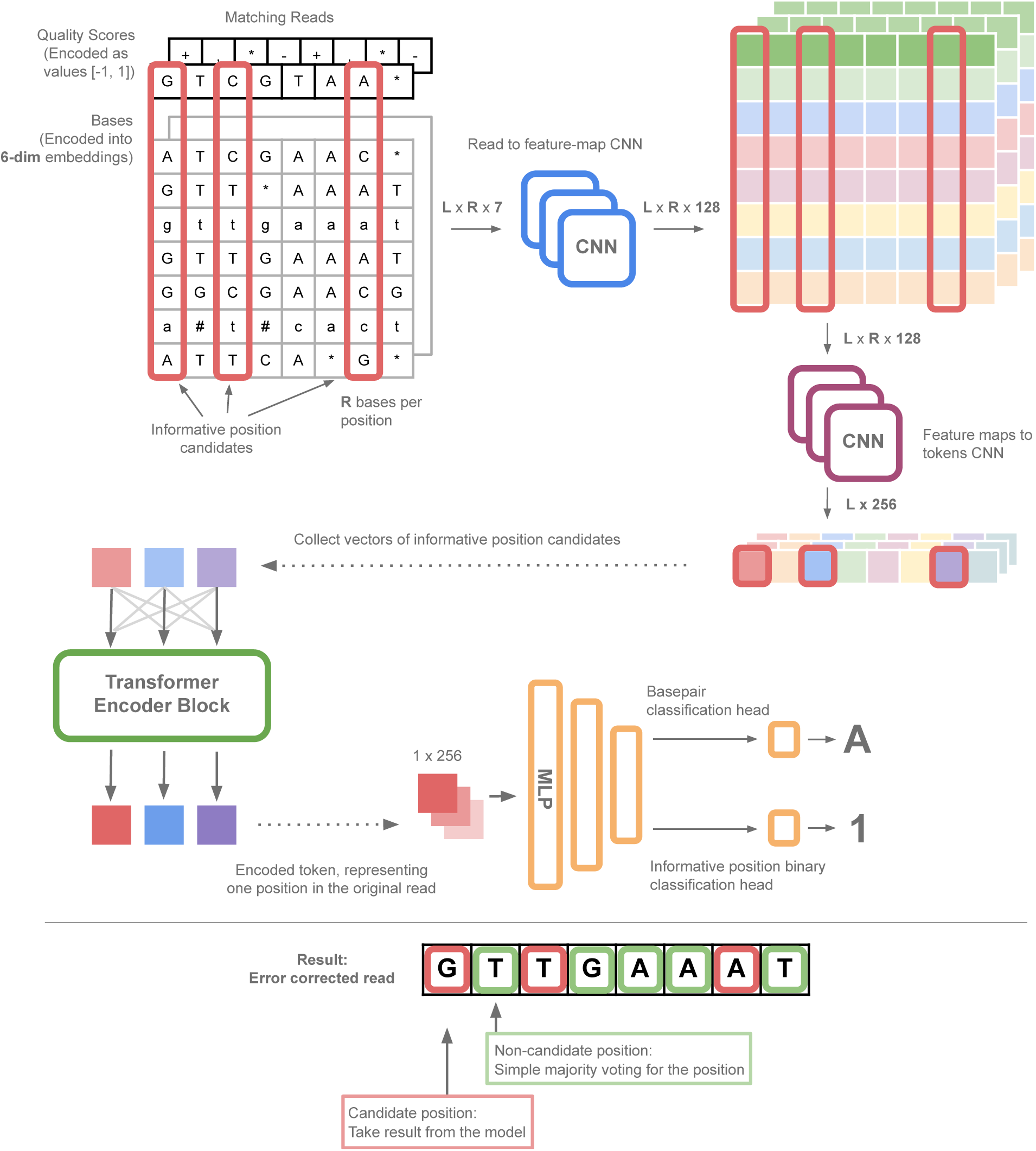
HERRO deep-learning model: The input to the model consists of embedded bases concatenated with base qualities, obtained by stacking pairwise alignments between a target read and the reads aligned to it. The first convolutional block extracts local information for each read independently. The second convolutional block aggregates information from different reads at a specific position in the target read. Next, only informative position candidates are indexed and fed into the Transformer encoder. The output from the Transformer encoder is used for base prediction. Bases at other target read positions are predicted using majority voting. The informative position prediction task is used for model training.

### HERRO effectively corrects ONT Simplex reads

We evaluated the alignment-based accuracy of the HERRO-corrected reads on several R10 and one R9 ONT datasets with existing high-quality assemblies. In terms of human samples, we considered HG002 data from Human Pangenome Reference Consortium (HPRC) and Genome In a Bottle (GIAB), CHM13 (pseudo-haploid) data from the Telomere-to-Telomere (T2T) consortium and I002C Indian male data with a recently released high-quality assembly (https://github.com/lbcb-sci/I002C or https://github.com/LHG-GG/I002C). We also used two inbred datasets *D. rerio* (zebrafish, TU strain) from the Vertebrate Genomes Project^22^ (VGP) and *A. thaliana* (thale cress, Col-0 strain) ^23^ (see Data Availability). The CHM13 data were sequenced using e8 chemistry and R9.4.1 flow cells. The other datasets were sequenced using e8.2 chemistry and R10.4.1 flow cells.

We aligned both uncorrected and corrected reads to the references using minimap2 and used bamConcordance tool^24^ to obtain a read quality measure (Q_concordance_ or Q_c_) and compute the rates of different types of errors (Methods, Analysis methods). Only primary alignments were considered in the evaluation. For reads successfully processed by HERRO, we observe that HERRO considerably improved Q_c_ for the majority of reads (Fig. 2a). There is a significant upward shift in the overall distribution of Q_c_ values for all considered datasets with an almost 100-fold increase in the median accuracy following correction (Fig. 2b). Extended Data Fig. 2 shows that a similar increase is also observed for higher quality (Q28) HG002 UL dataset.

**Fig. 2.**
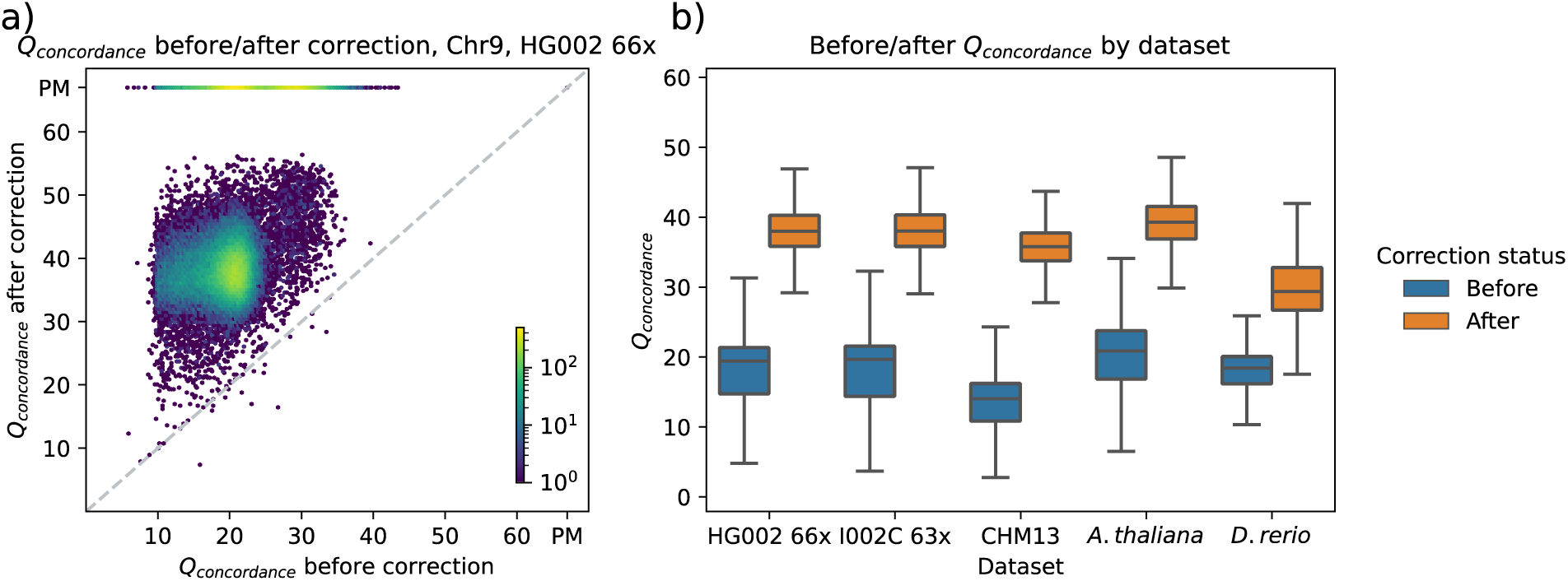
Qc before and after correction. a) A hexbin plot representing the empirical joint distribution of Qc values before and after correction for reads aligned to paternal and maternal chromosome 9 of the HG002 assembly v1.0.1^1,17,22–24^. The color indicates the density of data points in each hexagonal bin on a logarithmic scale. Bins above the dashed diagonal line indicate an improvement in accuracy after correction, while bins below it indicate a reduction. The “PM” at the end of each axis stands for “Perfect Match”. b) Box plot showing the distribution of Qc before and after correction across multiple datasets. I002C is a male Indian sample with a recently released reference (Data Availability). For all datasets, alignments that perfectly match the reference were excluded. Chromosomes of HG002 used to train the HERRO model were excluded (1, 2, 17, 18, and 20). CHM13 data was sequenced using R9.4.1 pore. All datasets shown here, except for A. thaliana, were sequenced with an ultra-long kit. Data in the box plots are displayed as follows: the centre line indicates the median, the bounds of the box represent the first and third quartiles (Q1 and Q3), and the whiskers extend to the minimum and maximum values within 1.5 times the interquartile range (IQR). Outliers beyond this range are plotted individually.

Table 1 shows the rates of different types of errors in the reads before and after correction. HERRO significantly reduced the rates across all error types, achieving, on average, a 50-fold reduction in the total number of errors across all datasets. Even for the *D. rerio* dataset, which showed the smallest reduction, the error rate has decreased nearly tenfold. The most substantial improvement was observed in mismatch errors, which in human datasets dropped from more than 100 errors per 10 kbp to less than 1. Similar reductions were observed for other types of errors except for non-homopolymer deletions in the *D. rerio* dataset. The higher error rate for non-HP deletions in this dataset may be due to the prevalence of highly repetitive, low-complexity elements in the genome^25^.

**Table 1.**
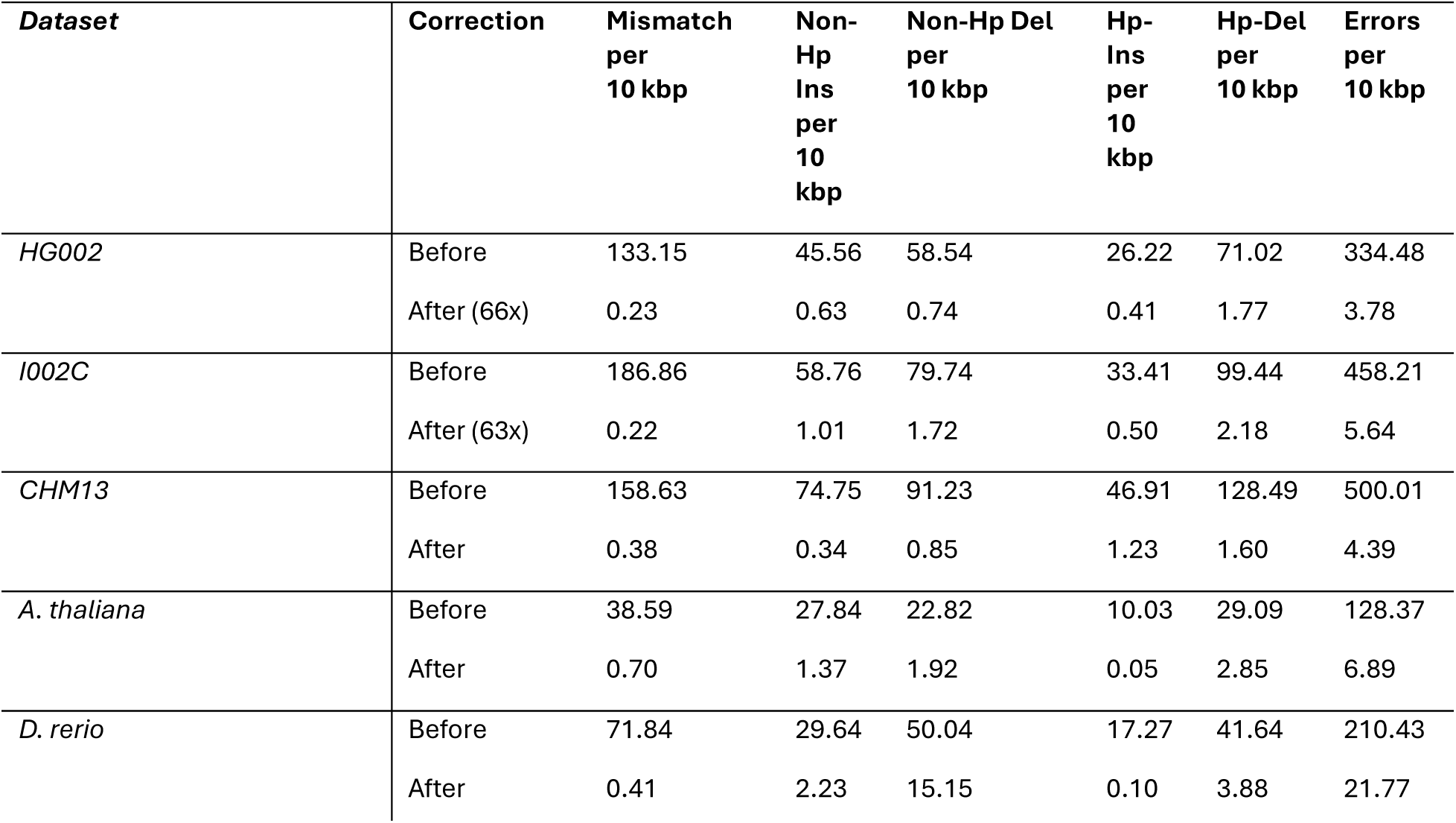
Counts of errors before and after correction, by type. We used bamConcordance to obtain counts of errors in reads before and after correction, based on alignments to a high-quality reference. In addition to the total number of errors, counts are provided for different categories: mismatch, non-homopolymer insertion, non-homopolymer deletion, homopolymer insertion, and homopolymer deletion.

### HERRO enables T2T-grade assemblies without high-accuracy long reads

Next, we evaluated the performance of HERRO-corrected ONT reads in a *de novo* assembly setting using several assemblers: Hifiasm, Verkko and LJA (only for datasets with very low heterozygosity). In addition to the datasets used to measure read accuracy in the previous section, we included three additional datasets: HG02818 from HPRC, HG005 from GIAB and a Drosophila melanogaster dataset (see Data Availability). All datasets were sequenced using e8.2 chemistry and R10.4.1 flow cells. For diploid human datasets, we also used available parental Illumina reads to enable the ‘trio analysis’ modes in Hifiasm and Verkko. HERRO-corrected reads were provided as high-accuracy long inputs, while original reads have been provided as ultra-long inputs. Corrected reads were downsampled to ∼35x for assembly if their coverages were considerably higher than 35x (38x or above) to control the effect of input coverage on assembly results and reduce computational requirements. To mitigate the issues related to contained read exclusion while running Hifiasm^26,27^, corrected reads were cut into 30 kbp subreads, filtering out any fragments shorter than 10 kbp. LJA only used the corrected reads as input. For details, see Methods. Key assembly metrics across selected assemblies are summarized in Table 2. To differentiate between assembly runs, the assembler name and input coverage used in correction are included in the run names. For more details regarding the different runs, refer to Supplementary Tables 1-8.

**Table 2.** Metrics of the assemblies for samples with a high-quality reference. We evaluated the assemblies of HERRO-corrected reads using high-quality references. NG50 and NGA50 metrics were calculated with minigraph^28^, while quality value (QV), switch errors, and Hamming errors were assessed using Merqury^29^. Gene completeness and duplication levels were measured with asmgene^21^ against CHM13 v2.0 for human samples and against high-quality assemblies for non-human samples. T2T contigs and scaffolds were identified using a script that searches for canonical telomeric motifs on the sides of assembled sequences and their alignment to a reference, with D. melanogaster being excluded due to its special telomere structure (Methods, Analysis methods). Sequences shorter than 500 kbp were excluded from the assembly analysis for A. thaliana (Supplementary Table 6). The HG002 assembly result corresponds to a 66x run (Supplementary Table 2), and the I002C assembly result corresponds to a 63x run (Supplementary Table 3). For comparison on the HG002 genome when HiFi + UL reads are used, we added results for an assembly from a recent publication^16^ by Cheng et al. They used an UL dataset, extracting only reads longer than 100kbp. For comparison on the HG002 genome when Duplex + UL reads are used, we added results evaluated for an assembly from a recent preprint^2^ by Koren et al. All assembly evaluations were done on scaffolds whenever scaffolds were produced.

| Assembly | Haplotype | Size (Gb) | NG50 (Mb) | NGA50 (Mbp) | QV | Hamming, switch errors (%) | Single-copy genes lost, duplicated, multi-copy genes lost (%) | Number of T2T contigs (plus scaffold) |
| --- | --- | --- | --- | --- | --- | --- | --- | --- |
| **Diploid assemblies** |  |  |  |  |  |  |  |  |
| HG002 (Verkko trio) | Maternal | 3.120 | 135 | 134 | 55.86 | 0.06,0.03 | 0.67,0.25, 5.34 | 9(13) |
|  | Paternal | 2.999 | 134 | 111 | 54.96 | 0.02,0.02 | 0.85,0.28, 8.03 | 9(12) |
| HG002 HiFi + UL (Verkko trio) Cheng et al<sup>16</sup> | Maternal | 3.026 | 112 | 110 | 56.50 | 0.07,0.04 | 0.91,0.21,5.29 | 4(5) |
|  | Paternal | 2.946 | 134 | 133 | 55.95 | 0.04,0.03 | 0.92,0.30,7.98 | 3(10) |
| HG002 Duplex + UL (Verkko trio) Koren et al<sup>2</sup> | Maternal | 3.037 | 147 | 114 | 53.59 | 0.05,0.03 | 0.78,0.34,5.19 | 6(11) |
|  | Paternal | 2.943 | 134 | 134 | 53.23 | 0.03,0.02 | 0.85,0.26,8.03 | 10(15) |
| I002C (Verkko trio) | Maternal | 3.057 | 146 | 124 | 61.06 | 0.29,0.27 | 0.58,0.27,4.34 | 6(13) |
|  | Paternal | 2.988 | 134 | 111 | 55.74 | 0.19,0.18 | 0.82,0.31,5.89 | 5(12) |
| **Low-heterozygosity assemblies** |  |  |  |  |  |  |  |  |
| CHM13 (LJA) | Pseudo-Haploid | 3.052 | 93 | 92 | 47.52 | N.A. | 0.04,0.009,0.55 | 4(4) |
| D. rerio (LJA) | Inbred | 1.451 | 56 | 56 | 53.26 | N.A. | 4.79,0.22,22.29 | 7(7) |
| A. thaliana (Hifiasm) | Inbred | 0.133 | 26 | 26 | 48.40 | N.A. | 0.05,0.02,0.87 | 3(3) |
| D. melanogaster (Hifiasm) | Inbred | 0.164 | 24 | 22 | 45.01 | N.A. | 0.0007,0.0006, 0.05 | N.A. |

#### Diploid human genomes

Multiple diploid human datasets, including HG002, I002C (Table 2), and HG02818 (Supplementary Table 1), were assembled. For each dataset, more than half of the 46 chromosomes were obtained as T2T contigs or scaffolds. Verkko generally achieves higher contiguity and produces more T2T chromosomes, whereas Hifiasm assemblies tend to have higher QV for some datasets (Supplementary Tables 1-4).

In multiple runs, the X and Y chromosomes were assembled as T2T contigs or scaffolds, with some runs remarkably achieving T2T contigs for both. While Hifiasm successfully assembled the X chromosome as T2T in some runs, only Verkko managed to assemble the challenging^30^ chromosome Y as T2T. These contigs were evaluated against HG002 assembly v1.0.1’s X and Y chromosomes using QUAST^31^ (Methods, Analysis methods), achieving genome fractions of 100% and 99.98% for X and Y, respectively, with a total of seven misassemblies (Table 3).

**Table 3.** Quast evaluation results of assembled T2T contigs of chromosomes X and Y. Assembled T2T contigs of chromosomes X and Y using corrected HG002 reads were evaluated against HG002 v1.0.1 using Quast. The reads were corrected at 66x coverage, assembled with Verkko, with and without using parental data, respectively. All the misassemblies are relocations.

| **Assembly** | **Genome fraction (%)** | **Misassemblies** | **Mismatches per 100kbp** | **Total mismatches** | **Indels per 100kbp** | **Total Indels** | **Indel <=5 bp** | **Indel >5bp** |
| --- | --- | --- | --- | --- | --- | --- | --- | --- |
| *ChrX, trio* | 100 | 2 | 0.19 | 286 | 8.43 | 13018 | 12672 | 346 |
| *ChrY, trio* | 99.98 | 5 | 1.18 | 736 | 4.08 | 2544 | 2410 | 134 |
| *ChrX, no-trio* | 100 | 2 | 0.17 | 270 | 8.43 | 13007 | 12656 | 351 |
| *ChrY, no-trio* | 99.98 | 5 | 1.16 | 724 | 4.07 | 2539 | 2405 | 134 |

We ran experiments at various coverage levels for each diploid ultra-long human dataset to evaluate the impact of different data coverages on correction and assembly. Most of these runs were performed on I002C data, which was not used for training and for which a high-quality reference is available. We observed a consistent increase in contiguity and the number of T2T chromosomes obtained as the data coverage increased. For instance, with 71x coverage (after filtering out short and low-quality reads), we achieved an NG50 of 146 Mbp for both haplotypes of I002C. At lower coverages, around 35x, the NG50 ranged from about 90 Mbp to 130 Mbp (Supplementary Tables 1-3).

To further isolate the effect of coverage used for correction on the quality of reads and, consequently, on the assembly, we compared I002C reads corrected at 45x, 63x and 71x coverage, which were then downsampled to 35x before being inputted as high-accuracy reads. Alongside these, the same uncorrected UL reads, fixed at 56x coverage were used as the UL input for Verkko (Supplementary Table 3). We observed an increasing trend in the number of T2T contigs or scaffolds as the coverage of the input reads to correction increased, with counts rising from 15 to 21 to 28. Due to a crash with the default version of Verkko (v2.0) during the 71x run, we used a newer version of Verkko for this experiment. To assess the effect of the version difference, we reran the assembly for the 45x run using the newer version of Verkko and found no change in the number of T2T chromosomes. Additionally, we calculated yak^14^ QVs for the I002C reads corrected at different input coverages, finding that QVs increased as coverage increased (Supplementary Table 9).

Besides HERRO-based runs, Table 2 also presents the evaluation of the HG002 trio Verkko assemblies from Cheng et al.^16^ and Koren et al.^2^, which correspondingly used HiFi and ONT Duplex data as high-accuracy inputs. The results are comparable, showing the power of correction of ultra-long ONT Simplex reads.

To test on non-UL human data, we corrected and assembled the HG005 dataset. Although this is not a UL dataset, it still has 10x coverage of reads 50 kbp or longer. In addition to the ONT reads, parental data was also provided to assemblers. When assembled with Verkko, 17 T2T chromosomes were obtained, including two that are T2T in single contigs. Full details about this run, as well as other runs for the HG005 dataset, can be found in Supplementary Table 4. Lastly, we corrected and assembled experimental higher-accuracy (Q28) ultra-long HG002 reads using a model trained on standard R10 data. Verkko produced 11 T2T contigs and 14 T2T scaffolds with the corrected reads but produced none without correction. Hifiasm, on the other hand, was able to produce 1 T2T contig and 18 T2T scaffolds without correction, and 7 T2T contigs and 12 T2T scaffolds with correction. The QV for each haplotype increased by around 10 with correction. These results demonstrate that error-correcting highly accurate data improves assembly contiguity, resulting in more T2T chromosomes in contigs with higher accuracy (Supplementary Table 10).

#### Difference between genomes

While evaluating error rates and assembly quality, we noticed discrepancies in the results depending on the reference genome used in the evaluation. Hence, we decided to quantify the differences in SNVs and indels (≤ 50 bp) across the different genomes.

Table 4 shows aggregated error rates of corrected reads evaluated by mapping to different high-quality assemblies. Since the error rate of corrected reads is very low, it can be almost neglected. The results show that differences in mean mismatch and indel rate are 0.15%– 0.19% and 0.14%–0.19%, respectively. In addition, we see notable differences in the NGA50 measure when evaluating genome assemblies against different references, with differences possibly exceeding 20% (Supplementary Table 12). It’s important to note that significant variations persist even when another haplotype from the same genome is used as a reference.

**Table 4.** The aggregated error rates of HERRO-corrected reads with respect to different assemblies. The error rates for HERRO-corrected reads were estimated by aligning them to several high-quality assemblies. All sets of reads were aligned to CHM13 v2.0 and CN1 ‘0.8 curated’^32^. HG002 reads were aligned to the HG002 v1.0.1 assembly, and I002C reads were aligned to the I002C v0.4 assembly. When aligning reads from different samples to the HG002 or I002C assemblies, only one autosomal haplotype was used (the paternal haplotype for HG002 and the maternal haplotype for I002C). Reads aligned to mitochondrial DNA or EBV sequences were excluded from the calculations. Indels longer than 50 bp were also excluded.

| **Corrected reads dataset** | **Reference** | **Mean error rate (%)** | **Mean mismatch rate (%)** | **Mean indel rate (%)** | **Median error rate (%)** | **Percentage mapped (%)** |
| --- | --- | --- | --- | --- | --- | --- |
| *HG002* | HG002 | 0.023 | 0.0019 | 0.021 | 0.014 | 99.995 |
|  | CHM13 | 0.30 | 0.16 | 0.14 | 0.16 | 99.94 |
|  | CN1 | 0.36 | 0.19 | 0.17 | 0.17 | 99.94 |
|  | I002C | 0.32 | 0.18 | 0.15 | 0.17 | 99.94 |
| *I002C* | HG002 | 0.34 | 0.18 | 0.16 | 0.17 | 99.97 |
|  | CHM13 | 0.33 | 0.18 | 0.15 | 0.17 | 99.97 |
|  | CN1 | 0.34 | 0.18 | 0.16 | 0.17 | 99.97 |
|  | I002C | 0.031 | 0.0056 | 0.026 | 0.014 | 99.97 |
| *CHM13* | HG002 | 0.31 | 0.15 | 0.16 | 0.16 | 99.98 |
|  | CHM13 | 0.048 | 0.0052 | 0.043 | 0.025 | 99.98 |
|  | CN1 | 0.36 | 0.17 | 0.19 | 0.17 | 99.98 |
|  | I002C | 0.33 | 0.16 | 0.16 | 0.17 | 99.98 |

These results highlight the importance of using high-quality assemblies and reads originating from the same genome when training deep-learning models. Since the error rate of mapping reads to a high-quality assembly from the same individual is low, we hypothesize that mapping highly accurate reads from one individual to a high-quality assembly of another can provide a rapid estimate of the differences between two genomes. Interestingly, the mean percentage of mismatches remains stable across genome pairs, offering an estimate of the number of single nucleotide variants (SNVs). The observed range of 0.15%-0.19% is higher than some previous estimates of 0.1%^33^. However, it is consistent with more recent findings where authors reported that the percentage of SNVs in segmental duplication regions is around 0.15%^34^. Moreover, a recently published preprint^35^ suggests similar differences between genomes for both SNVs and indels, with indels not having been quantified in previous studies.

#### Low-heterozygosity non-human genomes

We corrected and assembled ONT R10 data of highly homozygous samples for three non-human organisms: *A. thaliana (Col-0 strain)*, *D. melanogaster (iso-1 strain)*, and *D. rerio (*TU strain), with only *D. rerio* being ultra-long dataset. For *A. thaliana*, Hifiasm assembled three out of five chromosomes as T2T contigs (Table 2). LJA outperformed Hifiasm and Verkko in assembling *D. rerio*, achieving four T2T contigs compared to zero and one, respectively (Supplementary Table 5). Surprisingly, when we used *D. rerio* corrected reads longer than 90 kbp for assembly with LJA, the number of T2T contigs increased from four to seven (Supplementary Table 5).

#### R9.4.1 data

To test the ability of HERRO to correct R9.4.1 ONT Simplex reads, we corrected and assembled an ultra-long dataset for the CHM13^1^ genome. Across the different assemblers used, we achieved an NG50 of around 90 Mbp and up to five chromosomes as T2T contigs (Supplementary Table 8). Hifiasm and LJA produced similar results, with the most notable being that the Hifiasm assembly achieved the highest number of T2T chromosomes, while the LJA assembly produced more complete genes, as calculated using asmgene. Full results for all assembles can be found in Supplementary Table 8.

### Runtime assessment

Comprehensive runtime assessments for all datasets are presented in Supplementary Tables 1-8. For the HG002 dataset, with 66x input coverage, the correction process required approximately 69 hours of wall clock time in total. The majority of the time, about 64 hours, was spent running minimap2 to obtain all-vs-all alignments, using 64 threads and peaking at 102 GB of RAM. Following this, the correction phase took approximately 5 hours with peak RAM consumption of 550 GB using 64 threads. More details can be found in the Methods.

## Discussion

Long high-accuracy PacBio HiFi and ONT Duplex reads have created new opportunities for reconstructing large and complex genomes. Their low error rates have enabled the assembly of challenging regions, such as segmental duplications, by distinguishing tiny variations between each segment. Moreover, these reads and new assembly methods considerably improved haplotype resolution. However, to reconstruct the most complex regions, like centromeres and other satellite arrays, and to fully phase the haplotypes, ultra-long ONT reads are required. Due to the higher error rate, their usage as a basis for the construction of high-resolution assembly graphs was limited, leading to a hybrid strategy in which an assembly graph is first built from long high-accuracy reads, with further integration of ultra-long ONT data. This approach combined with trio analysis and manual curation, has led to several breakthroughs. These include the telomere-to-telomere (T2T) reconstruction of a pseudo-haploid human genome, the reconstruction of complete human chromosome Y, and the T2T haplotype-resolved assembly of the HG002 diploid human genome.

We began our work with the goal of fully leveraging the potential of ultra-long ONT Simplex reads, thereby removing the need for PacBio HiFi or ONT Duplex reads in assembly projects. To achieve this, we decided to try reducing the error rate of ONT reads through self-correction. However, heuristics-based approaches, such as those used in Racon^36^, PECAT^13^, Hifiasm^14^, or HiCanu^37^, have proven inadequate as they may overcorrect important, informative positions that are crucial for reconstructing segmental duplications and phasing polyploid genomes. Therefore, we set out to develop a new error-correcting method focused on those key positions.

Using HERRO, a deep learning model trained on ONT Simplex reads labelled with the high-quality HG002 assembly, allows us to correct reads to a level where they can be used in existing assemblers as long accurate reads, without overcorrecting informative positions. However, these assemblers were not initially designed to fully exploit the potential of ultra-long accurate reads. Inspired by RAFT^26^, we chopped the corrected UL reads before inputting them into Hifiasm, which improved the results. In contrast, our tests showed that using unchopped reads resulted in lower-quality genomes produced by Hifiasm. For Verkko and LJA, chopping reads was unnecessary.

The results indicate that we can achieve high contiguity comparable to approaches that combine HiFi or Duplex with ultra-long ONT reads. However, we believe the current assembly results are not yet optimal due to several factors, including potential under- or over-correction by HERRO and the need for assemblers to be further tailored for ultra-long reads. Notably, we demonstrate routine reconstruction of complete human chromosomes X and Y from ultra-long ONT reads. This gives us hope that with further improvements in error correction, and the optimization of assemblers for high-accuracy ultra-long reads, we will be able to routinely assemble even the most complex parts of the genomes (e.g. rDNAs) and fully assemble polyploids, achieving chromosomes in T2T contigs, without reliance on high-accuracy long reads.

We acknowledge that there is room for improvement in all parts of the pipeline, including better alignment (e.g., using optimized pairwise aligners such as Edlib^38^ or A*^39^), reducing memory and computational requirements, optimizing window length, and gaining a better understanding of the remaining errors. Using minimap2 to find all-vs-all alignments may be suboptimal in terms of both runtime and accuracy. Modern assemblers like Hifiasm and Verkko use built-in overlappers specifically designed for long, high-accuracy (or ultra-long) reads. Developing an overlapper tailored specifically for ONT pre-corrected reads could produce more accurate overlaps in less time. Some promising approaches involve parallelizing specific components of minimap2 on CPU^40^ or GPU^41^ or developing entirely new algorithms.

Furthermore, HERRO handles indels up to 30 nucleotides long, and exploring options for longer indels is important. Training on more data, including other organisms, could lead to even lower error rates. Another area worth exploring is the assembly of metagenomic samples^42–44^. Moreover, here we haven’t yet explored the impact of highly accurate long ONT reads on the assemblers tailored to produce collapsed haploid assemblies such as Raven^45^ and Flye^9^. While these assemblers do not reconstruct individual haplotypes, they require neither UL reads nor parental (or other long-range) data, while still providing meaningful assemblies often suitable as references for population studies. Increased accuracy of the input data allows for adjustments throughout the assembly pipeline, improving performance and separation of more diverged segmental duplication copies. Finally, adding methylation information (e.g., using Rockfish^46^ or similar tools) might improve the identification of informative positions.

Our results indicate that mapping long, highly accurate reads to a high-quality assembly from another individual results in differences of 0.15%–0.19% in mismatches and 0.14%– 0.19% in short indels. Additionally, we found that the structural concordance metrics, such as NGA50, can vary depending on the reference genome used for assessment. These observations highlight the importance of using de novo assemblies generated from reads from the same individual rather than relying on reference sequences built from a different individual. While we acknowledge that our sample size is small and that a more comprehensive analysis is needed, this demonstrates the potential of using long, highly accurate reads to estimate genomic differences quickly. These insights may provide guidelines for future machine learning methods that rely on read mapping, such as error correction, DNA modification detection, and similar.

In summary, this paper introduces HERRO, an error correction method that enables high-quality assembly using ONT Simplex reads as the sole long-read sequencing technology. This approach results in lower costs and reduced genomic DNA requirements while achieving state-of-the-art assemblies. Moreover, by providing a model that supports R9.4.1 reads, HERRO allows researchers to use even older data in state-of-the-art assembly pipelines.

Looking ahead, accurate long and ultra-long ONT reads open new possibilities for studying more complex genomes, including those with different ploidy and aneuploidy levels.

## Methods

### Data preprocessing

Data preprocessing consists of read basecalling, adapter trimming, read splitting on middle adapters and filtering out short and low-quality reads. All datasets were basecalled using Dorado (https://github.com/nanoporetech/dorado), with the model version depending on the pore type. R9.4.1 data was basecalled using the dna_r9.4.1_e8_sup@v3.6 model, R10.4.1 4 kHz data using the dna_r10.4.1_e8.2_400bps_sup@v4.1.0 model and R10.4.1 5 kHz data using the dna_r10.4.1_e8.2_400bps_sup@v4.3.0 model. Adapter trimming is done using Porechop^47^, wherein we recognize and remove adapter sets with at least one match over 95%. Next, we detect middle adapters and split reads using the *split_on_adapter* script from Duplex Tools (https://github.com/nanoporetech/duplex-tools). Dorado versions v0.5.0 or higher implement adapter trimming and splitting, making Porechop and Duplex Tools unnecessary. Finally, reads shorter than 10 kbp and those with a mean base quality below 10 are discarded using SeqKit^48,49^. Only the resulting reads will be used in the error correction process. Read filtering by length and quality was used in our experiments and is recommended but is not built into the main pipeline. Additional details and the preprocessing script are available in the Supplementary Information.

### Feature generation

To generate features, we first perform all-vs-all alignment using minimap2^21,50^. If multiple alignments between two reads exist, we keep only the first one, assuming that alignment is optimal. We perform error correction on each read independently. The read that will be corrected is named *target* read, while the reads aligned with it are named *query* reads.

Initially, we divide a target read into chunks by defining non-overlapping windows of length *L*_*T*_ = 4096 base pairs (bp). We extract the corresponding segments from the query reads for each window based on the alignments (CIGAR string). Any query segment that doesn’t cover the entire window is discarded. Moreover, a query segment for a window is also discarded if any insertion or deletion is longer than 30 bp. Next, we sort segments by alignment accuracy and retain only the top 30 segments. All segments are retained if there are fewer than 30 for a window. For each window, we define two tensors: *B* ∈ ℝ^31^ ^×^ ^*L*^ and *Q* ∈ ℝ^31^ ^×^ ^*L*^, corresponding to the stacked pairwise bases and stacked pairwise base qualities, respectively. The first row in both tensors corresponds to the target read, while the subsequent rows contain the sorted segments in decreasing order. If fewer than 30 segments exist, we insert a special “no alignment” token (’.’) in the remaining rows.

Moreover, we define a special gap token (’*’) to denote insertions with respect to the target read. This results in a total tensor length 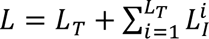 where 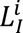 is the length of the longest insertion at the target position *i*. Each column, which can be either a target or inserted position, is later called independently. Information about the alignment strand is given implicitly; bases from different strands are encoded differently. There are 11 tokens: five bases (A, C, G, T, and ‘*’) for each strand (forward and reverse) plus a no-alignment token. Base qualities for the insertions are set to 0.

In addition to features, we also extract information about informative position candidates. We define a position as an *informative position candidate* if at least two different bases (including gap) are supported by a minimum of three reads at that specific position. Positions that do not meet this criterion are considered to be *non-candidate positions*. The deep learning model predicts only the target bases for informative position candidates.

### Deep learning model and training

#### Model architecture

HERRO deep learning model consists of three parts. In the first part, we update information for each base based on its local neighbourhood before pooling information from each base at the same position to obtain a vector representation for each position. Initially, the base tensor is embedded using an embedding layer, resulting in the embedded bases *E*_*B*_ ∈ ℝ^31×*L*×6^. This tensor and the quality base tensor are concatenated into an input tensor *I* ∈ ℝ^31×*L*×7^, processed through two convolutional blocks. Each convolutional block consists of 1D convolution, Batch Normalization and Rectified Linear Unit (ReLU). The output of the first stage is the intermediate tensor *H*(1) ∈ ℝ^*L*×256^. In the second part, we focus only on representations from informative position candidates, updating them based on interactions with other candidates. By concentrating only on the candidates, we process only a fraction of all positions, drastically reducing the computational demands. The tensor *H*(2) ∈ *H*^*S*×256^, containing representations of informative position candidates are fed into a Pre-LN Transformer encoder^51,52^. The output from the encoder is passed to the final part of the model, which consists of two classification heads: the base prediction head and the informative position prediction head. The base prediction head consists of a fully connected layer followed by a softmax layer, produces base probabilities *Y*_*B*_ ∈ *H*^*S*×5^ for every candidate position. For each candidate position, it predicts whether the correct base is one of the canonical bases (A, C, G, T) or a gap symbol (’*’). The informative position prediction head, defined as a fully connected layer followed by a sigmoid, outputs the probabilities *Y*_*I*_ ∈ *H*^*S*×1^, which denotes whether each candidate is an informative position. A candidate position is deemed *informative* if different haplotypes have different bases at that position. The block diagram, which displays the entire architecture with defined parameters, is shown in Extended Data Fig. 3.

#### Model training

We perform multi-task learning to find the optimal model’s parameters. The total model loss for one window is the sum of the losses from each candidate position. The total loss for the candidate position *i*, where *i* ∈ {1, 2, …, *S*}, is a sum of two losses: base prediction loss and informative position prediction loss and is given by:

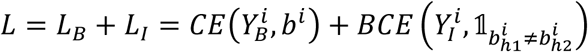

where *CE* is the cross-entropy loss, *BCE* is the binary cross-entropy loss, 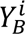 represents the predicted base probabilities, *b*^*i*^ is the true base, 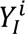 denotes the predicted probability that a candidate position is informative, and 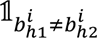 is the indicator function that equals to 1 if the bases on haplotypes ℎ_1_ and ℎ_2_ are different, and 0 otherwise.

We trained the model on four A100 GPUs for a maximum of one million steps. The batch size was 128, so the model saw 128 million examples. Validation was performed every 2000 steps, and we selected the model with the highest validation F1 score for the base prediction task. The Adam optimizer was used for training with default β_1_ = 0.9 and β_2_ = 0.999 parameters. We used a cosine learning rate scheduler with an initial learning rate of 3 × 10^−4^. Weight decay was set to 0.1 and gradients were clipped using the ℓ^2^-norm with a maximum norm value of 1. The dropout value in transformer was set to 0.1. The complete list of hyperparameters can be found in Supplementary Table 11.

#### Training dataset

We used a portion of the HG002 dataset internally sequenced by ONT for training (HG002 internal), specifically chromosomes 1-2 and 17-18. Chromosome 20 was used for validation. The remaining chromosomes were excluded from training and model selection. The reads were first aligned to the recently published high-quality assembly using minimap2. This diploid assembly allowed us to assign the chromosome and haplotype to each read. Only reads with primary alignments to the specified chromosomes and mapping quality of 2 or higher were processed further, as previously described. To identify correct bases and informative positions, we mapped both contigs (haplotypes) corresponding to the assigned chromosome to each read. We extracted the correct base from the assigned haplotype for each position on a read. If the bases differed between the two haplotypes at a particular position, that position was labelled informative. If there was no mapping for the non-assigned haplotype, all positions were automatically labelled as non-informative.

### Consensus

After obtaining predictions from the deep learning model for the candidate positions, we determine a consensus sequence for each window. For non-candidate positions, we use simple majority voting. If two or more bases (including gaps) occur with the same frequency and one of them is a target base, we copy the target base; otherwise, the base is chosen randomly. For the candidate positions, we select the base with the highest predicted probability. Consecutive windows are stitched into a single sequence. If a window contains fewer than two alignments, the window is discarded, and the read is split. The resulting sequences are written to a FASTA file.

## Analysis methods

### Generating assemblies with Hifiasm

We ran Hifiasm v0.19.8 to generate assemblies for all corrected datasets, subsampling them to 35x coverage if they exceeded 38x. The corrected reads were chopped to 30 kbp and filtered to retain only those longer than 10 kbp. According to our experiments, this was necessary because Hifiasm did not perform optimally with longer reads as high-accuracy inputs. This could be related to the nature of its algorithm^26,27^. Reads that were trimmed (using porechop and duplex_tools on the fastq or trimmed during basecalling by Dorado v0.5.0 and above) but not yet corrected and that were 50 kbp or longer were used as the UL input for Hifiasm. In the case of haploid genomes, primary assemblies are evaluated. For diploid samples, parental data were processed using yak and used with Hifiasm’s built-in trio-binning mode to obtain phased assemblies. For detailed commands, versions and parameters, see Supplementary Information, Assembly section.

### Generating assemblies with Verkko

Verkko was run on all corrected datasets to generate assemblies, subsampling them to 35x if their coverage was 38x or above. We generally used Verkko v2.0 by default; however, in some instances where it crashed, we used a newer version (Supplementary Information, Assembly). For Verkko, the corrected reads were used as is, without chopping. Trimmed, uncorrected reads were used as the UL input for Verkko without filtering by length. For diploid samples, Meryl and Merqury were used to process parental data in Verkko’s trio-mode to generate phased assemblies. Homozygous or haploid samples were run with the “--haploid” option to improve the alignment of ONT reads. Refer to the Supplementary Information, Assembly section for detailed commands, versions and parameters.

### Generating haploid assemblies with LJA

LJA (v0.2) was run on all homozygous or haploid corrected datasets to generate assemblies, subsampling them to 35x if they were 38x or above. Only the downsampled corrected reads were used as input for LJA. For detailed commands, see Supplementary Information, Assembly section.

### Evaluation of gene completeness

We evaluated the gene completeness of assemblies using three different methods – asmgene, BUSCO^53,54^ and compleasm^55^.

For asmgene, a reference assembly was used for each sample. Minimap2 2.26-r1175 was used to align cDNA sequences to the reference assembly and then to the constructed assembly (for diploid samples, each haplotype’s assembly is assessed separately). We then ran asmgene from paftools 2.26-r1175 and used its outputs to calculate how many single-copy or multi-copy genes found on the reference remained single- or multi-copy on the assembly. From this, we calculated the proportion of single-copy genes on the reference that were no longer single-copy on the assembly, the proportion of single-copy genes that became duplicated, and the proportion of multi-copy genes that were no longer multi-copy by the following:

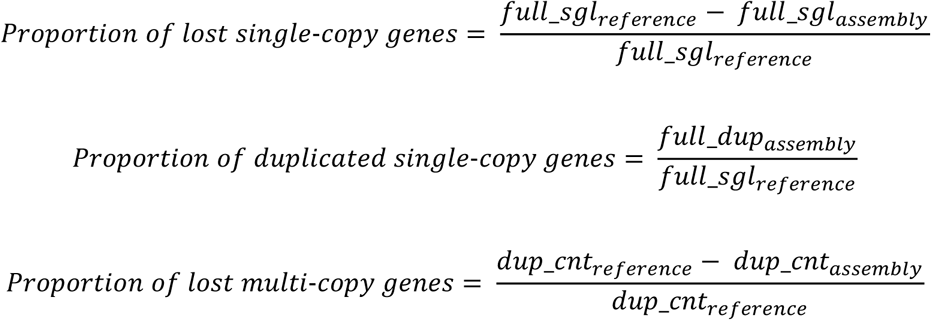

For non-human datasets, we also additionally ran BUSCO 5.6.1 and compleasm 0.2.5 to obtain a measure of gene completeness by checking for the existence of conserved single-copy orthologs in the assembly. For detailed commands and reference for each sample, see Supplementary Information, Evaluation.

### Reference-based evaluation of assemblies

Minigraph 0.20-r574-dirty with paftools 2.26-r1175 was used to assess each assembly’s contiguity, completeness and correctness against a reference by alignment. We first aligned assembly sequences to the reference sequences using minigraph and indexed the reference with Samtools^56^ 1.19.2 (Using htslib^57^ 1.19.1). Paftools then used this information to calculate metrics such as NG50 and NGA50. NG50 is calculated by summing the lengths of the assembled sequences in descending order, and NGA50 is similar but uses aligned segments of assembly sequences to the reference; both metrics identify the sequence length at the point where cumulative length equals or exceeds 50% of the genome length. The percentage of the reference sequences aligned to a sequence in the assemblies measures assembly completeness (Rcov; Supplementary Tables 1-8, Supplementary Table 10). Additionally, Quast 5.2.0 with minimap2 2.24-r1122 was used to evaluate T2T X and Y contigs for HG002 against the X and Y chromosomes in HG002 assembly v1.0.1. For detailed commands and the references used, see Supplementary Information, Evaluation.

### K-mers based evaluation of assemblies

We used Merqury (with Meryl) and yak to assess assemblies using short, accurate reads from the same sample. k-mers in the short reads were counted and compared with the k-mers in assembly sequences to estimate the assembly’s QV (a log-scaled measure of base error rate). For diploid samples, maternal- and paternal-specific k-mers were derived from the short reads of the parents of the assembled sample. This approach identifies k-mers from both parents in the assembled sequences, allowing it to identify switch and Hamming errors. Refer to the Supplementary information, Evaluation for detailed versions, commands and parameters.

### Counting T2T chromosomes in assemblies

The script from https://github.com/prasad693/Tel_Sequences was used to identify telomere-to-telomere (T2T) sequences in the assemblies of all samples, except for D. melanogaster, for which telomeres do not consist of short motif repeats^58^. The script searches for species-specific canonical telomeric motifs at the ends of each sequence in the assembly and attempts to align these sequences to a reference sequence. Assembly sequences that cover the full length of the reference chromosome and also have motif repeat counts exceeding a specified threshold were considered T2T. For detailed commands, see Supplementary Information, Evaluation.

### Measurement of reads’ quality

Reads before and after correction were aligned to a reference for samples with a high-quality reference using minimap2 2.26-r1175. The bamConcordance tool was used on the alignments to calculate Q_concordance_ (Q_c_), a score that is a log-scaled measure of base error probabilities. It also outputs the counts of different types of errors – mismatch, non-homopolymer insertion, non-homopolymer deletion, homopolymer insertion, and homopolymer deletion of the reads compared to the reference. Only the records for the primary alignments were used. Yak 0.1-r69-dirty was used to estimate QV, a log-scaled measure of base error probabilities of the reads for I002C reads corrected at different coverages. Corrected reads from HG002 (at 66x), I002C (at 63x) and CHM13 (67x) were also aligned to different high-quality human assemblies to estimate the error rates of reads with respect to them. Error rates were calculated by the following:

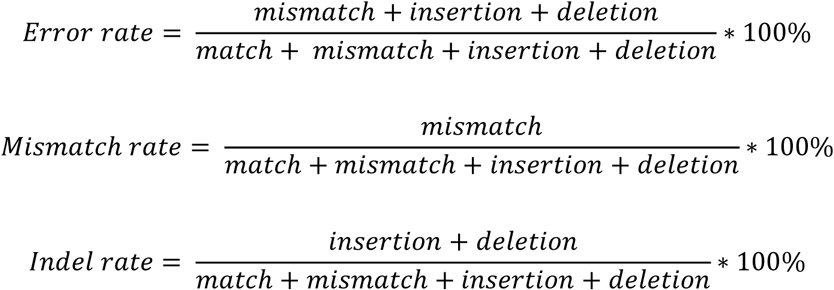

Indels longer than 50 bp are typically classified as structural variants (SVs) and were therefore excluded from the analysis. Additionally, reads aligned to mitochondrial DNA or EBV were also excluded. For detailed commands and alignment references, see Supplementary Information, Evaluation.

### Measurement of computational resources used

The elapsed time, CPU time, and peak RAM usage were measured for the all-versus-all read alignment, correction, and assembly steps. These numbers were taken from the output log for Hifiasm assembly runs and obtained with “/usr/bin/time -v” for the rest. All-versus-all alignment was performed on the nodes of a computing cluster, using 64 threads on an AMD EPYC 7543 32-core processor for each run. GPU inference (correction step) was run on GPU nodes on a cluster, with 4 NVIDIA A100-SXM4-80GB being used for each run, along with 64 worker threads on an AMD EPYC 7713P 64-core processor. Most of the assembly runs used 64 threads on an AMD EPYC 7742 64-core processor, except the run ‘Danio rerio, 80x LJA’ in Supplementary Table 5 which used 16 threads.

## Data availability

HG02818 ULONT data: https://s3-us-west-2.amazonaws.com/human-pangenomics/index.html?prefix=submissions/3471c92c-5b6a-4c20-9953-13e3a851debc--UCSC_HPRC_PLUS_nanopore_WashU_samples/HG02818/ ; HG02818 and parents’ short reads data: https://s3-us-west-2.amazonaws.com/human-pangenomics/index.html?prefix=working/HPRC_PLUS/HG02818/raw_data/Illumina/ ; HG005 ONT data: https://s3-us-west-2.amazonaws.com/human-pangenomics/index.html?prefix=submissions/5b73fa0e-658a-4248-b2b8-cd16155bc157--UCSC_GIAB_R1041_nanopore/HG005_R1041_Sheared/ ; HG005 and parents’ short reads data: https://s3-us-west-2.amazonaws.com/human-pangenomics/index.html?prefix=working/HPRC_PLUS/HG005/raw_data/Illumina/ ; HG002 ULONT data: https://s3-us-west-2.amazonaws.com/human-pangenomics/index.html?prefix=submissions/5b73fa0e-658a-4248-b2b8-cd16155bc157--UCSC_GIAB_R1041_nanopore/HG002_R1041_UL/dorado/v0.4.0_wMods/ (Only the data from the four files with ‘ULCIR’ or ‘ULNEB’ in their filenames were used) ; HG002 and parents’ short reads data: https://s3-us-west-2.amazonaws.com/human-pangenomics/index.html?prefix=working/HPRC_PLUS/HG002/raw_data/Illumina/ ; D. melanogaster ONT data: https://labs.epi2me.io/open-data-drosophila/ ; D. melanogaster short reads data: https://www.ncbi.nlm.nih.gov/sra/SRR6702604 ; A.thaliana ONT data: https://www.ncbi.nlm.nih.gov/sra/?term=SRR29061597 (See Supplementary Information, Additional notes on Arabidopsis thaliana data); A. thaliana Illumina data: https://ngdc.cncb.ac.cn/gsa/browse/CRA005350 ; D. rerio ULONT data: https://genomeark.s3.amazonaws.com/index.html?prefix=species/Danio_rerio/fDanRer17/genomic_data/ont/ (PAG68679 was not used for consistency because it has a different data sampling frequency from all the rest) ; D. rerio short reads data: https://www.ncbi.nlm.nih.gov/sra?linkname=bioproject_sra_all&from_uid=1029986 ; CHM13 ULONT data: https://s3-us-west-2.amazonaws.com/human-pangenomics/index.html?prefix=T2T/CHM13/nanopore/fast5/ (some partitions were skipped due to reasons such as having too many fast5 files, see Supplementary information: CHM13 data partitions used); CHM13 short reads data: https://www.ncbi.nlm.nih.gov/sra/?term=SRX1082031 ; HG002 v1.0.1: https://github.com/marbl/HG002 ; CHM13 v2.0: https://github.com/marbl/CHM13 ; I002C v.04: https://github.com/lbcb-sci/I002C or https://github.com/LHG-GG/I002C; CN1: https://github.com/T2T-CN1/CN1 ; D. rerio assembly used as reference: https://www.ncbi.nlm.nih.gov/datasets/genome/GCA_033170195.2/ ; A. thaliana assembly used as reference: https://ngdc.cncb.ac.cn/gwh/Assembly/21820/show ; D. melanogaster assembly used as reference: https://www.ncbi.nlm.nih.gov/datasets/genome/GCA_018904365.1/ ; Human cDNA used: https://ftp.ensembl.org/pub/release-111/fasta/homo_sapiens/cdna/Homo_sapiens.GRCh38.cdna.all.fa.gz ; D. melanogaster cDNA used: https://ftp.ensembl.org/pub/release-111/fasta/drosophila_melanogaster/cdna/Drosophila_melanogaster.BDGP6.46.cdna.all.fa.gz ; A. thaliana cDNA used: https://ftp.ensemblgenomes.ebi.ac.uk/pub/plants/release-58/fasta/arabidopsis_thaliana/cdna/Arabidopsis_thaliana.TAIR10.cdna.all.fa.gz ; D. rerio cDNA used: https://ftp.ensembl.org/pub/release-111/fasta/danio_rerio/cdna/Danio_rerio.GRCz11.cdna.all.fa.gz; HG002 UL Q28 dataset used: https://labs.epi2me.io/gm24385_ncm23_preview/; HG002 data (HG002 internal) used for training R10 model will be available for peer-reviewers upon request; Data for I002C will be available prior to publication; HG002 data used for training R9 model: https://labs.epi2me.io/gm24385_2021.05/ ; HiFi reads’ assembly in Table 2: https://s3-us-west-2.amazonaws.com/human-pangenomics/index.html?prefix=submissions/53FEE631-4264-4627-8FB6-09D7364F4D3B--ASM-COMP/HG002/assemblies/verkko_1.3.1/trio/ ; Duplex reads’ assembly in Table 2: https://obj.umiacs.umd.edu/marbl_publications/duplex/HG002/asms/duplex_50x_30xUL_t rio.tar.gz ; Most of the assemblies evaluated in this paper: https://zenodo.org/records/13702708 ; Q28 ULONT assemblies evaluated in this paper: https://zenodo.org/records/13703130 ;

## Code availability

HERRO code is available at https://github.com/lbcb-sci/HERRO. The pre-trained models are available at https://zenodo.org/records/12683277.

## Acknowledgements

This work has been jointly funded by AI Singapore and Oxford Nanopore Technologies under the grant AISG2-100E-2021-076. D. S. has also been funded by ARAP A*STAR scholarship, National Medical Research Council, Singapore, under the grant MOH-000649-01 (Rapid diagnostic of infectious diseases based on nanopore sequencing and AI methods) and Croatian Science Foundation under the grant DOK-2018-09-8714.

We would like to acknowledge Martin Frederik Schmitz for his invaluable help in drawing images.

The achievements detailed in this manuscript would not have been possible without the tremendous efforts of the scientific community. Their production of high-quality references and provision of open access to sequenced reads were instrumental in this work.

We would like to thank the Human Pangenome Reference Consortium (HPRC), the Telomere-to-Telomere (T2T) Consortium, the Vertebrate Genomes Project (VGP) and the Genome in a Bottle (GIAB) Consortium for making sequencing datasets publicly available. We thank George Dodd of Oxford Nanopore Technologies (ONT) for providing the Arabidopsis thaliana dataset. We thank Sean McKenzie for his assistance with datasets and Cheng Yong Tham for assistance in transferring data. We would like to acknowledge the National Genome Research Institute (NHGRI) for funding the following grants which are in support of creating the human pangenome reference: 1U41HG010972, 1U01HG010971, 1U01HG010961, 1U01HG010973, 1U01HG010963, and the Human Pangenome Reference Consortium (https://humanpangenome.org/).

## Author contributions

M. Š. and P. F. S. conceived the project. D. S. designed and implemented HERRO, with contributions from M. Š., S. N. and D. L.; D. L. performed the assembly experiments and bioinformatics analysis, assisted by M. Š., D. S. and S. N.; D. S., D. L., and M. Š. organized the manuscript. D. L., D. S., and M. Š. wrote the manuscript, with additional input from P. F. S. and S. N.; M. Š. and P. F. S. supervised the project, while M. Š. and S. N. provided mentorship and support throughout the project.

---

**Extended Data Fig. 1.**
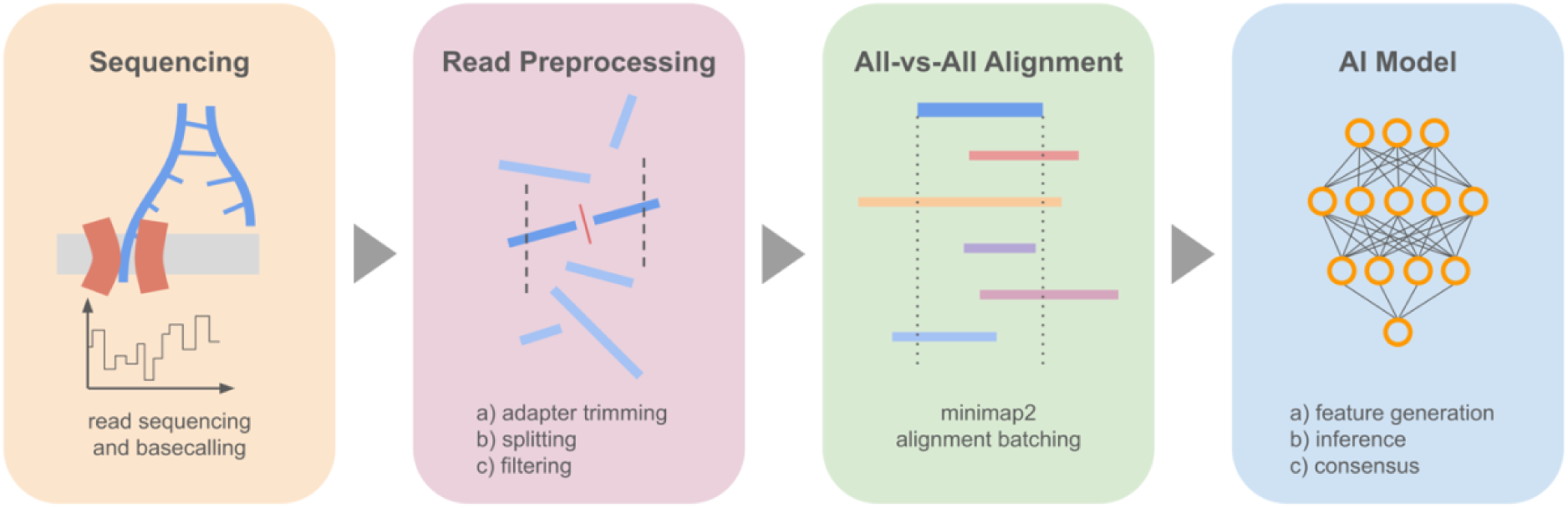
Overview of the HERRO method pipeline: The input to the pipeline are ONT sequencing reads. After trimming adapters and splitting at middle adapters, short and low-quality reads are filtered out. Next, all-vs-all alignments are performed using minimap2. In feature generation, we chunk the target read and its alignments into multiple windows. For each window, we define two matrices representing stacked bases and base qualities. These windows are then processed by a deep-learning model. To predict the correct base for the candidate positions, we use predictions from the model. For non-candidate positions, we apply simple majority voting. The final error-corrected read is obtained by stitching together the consensus sequence from each window.

**Extended Data Fig. 2.**
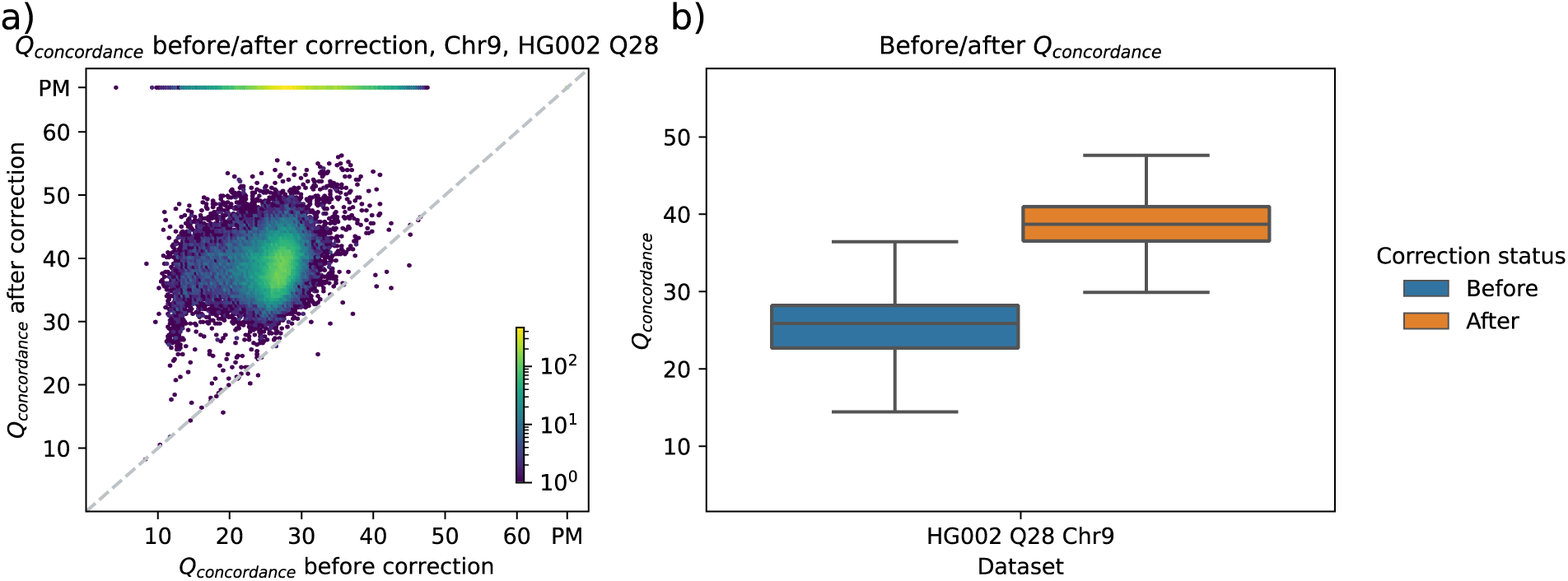
Qc before and after correction for Q28 ultra-long HG002 reads. a) A hexbin plot representing the data density for specific pairs of Qc values before and after correction for reads aligned to paternal and maternal chromosome 9 of the HG002 assembly v1.0.1. The color intensity indicates the density of data points in each hexagonal bin on a logarithmic scale. Bins above the dashed diagonal line indicate an improvement in accuracy after correction, while bins below it denote a reduction. The “PM” at the end of each axis stands for “Perfect Match”. b) This plot shows the distribution of Qc before and after correction. Alignments that perfectly match the reference were excluded. Data in the box plots are presented as follows: the centre line indicates the median, the bounds of the box represent the first and third quartiles (Q1 and Q3), and the whiskers extend to the minimum and maximum values within 1.5 times the interquartile range (IQR). Outliers beyond this range are plotted individually.

**Extended Data Fig. 3.**
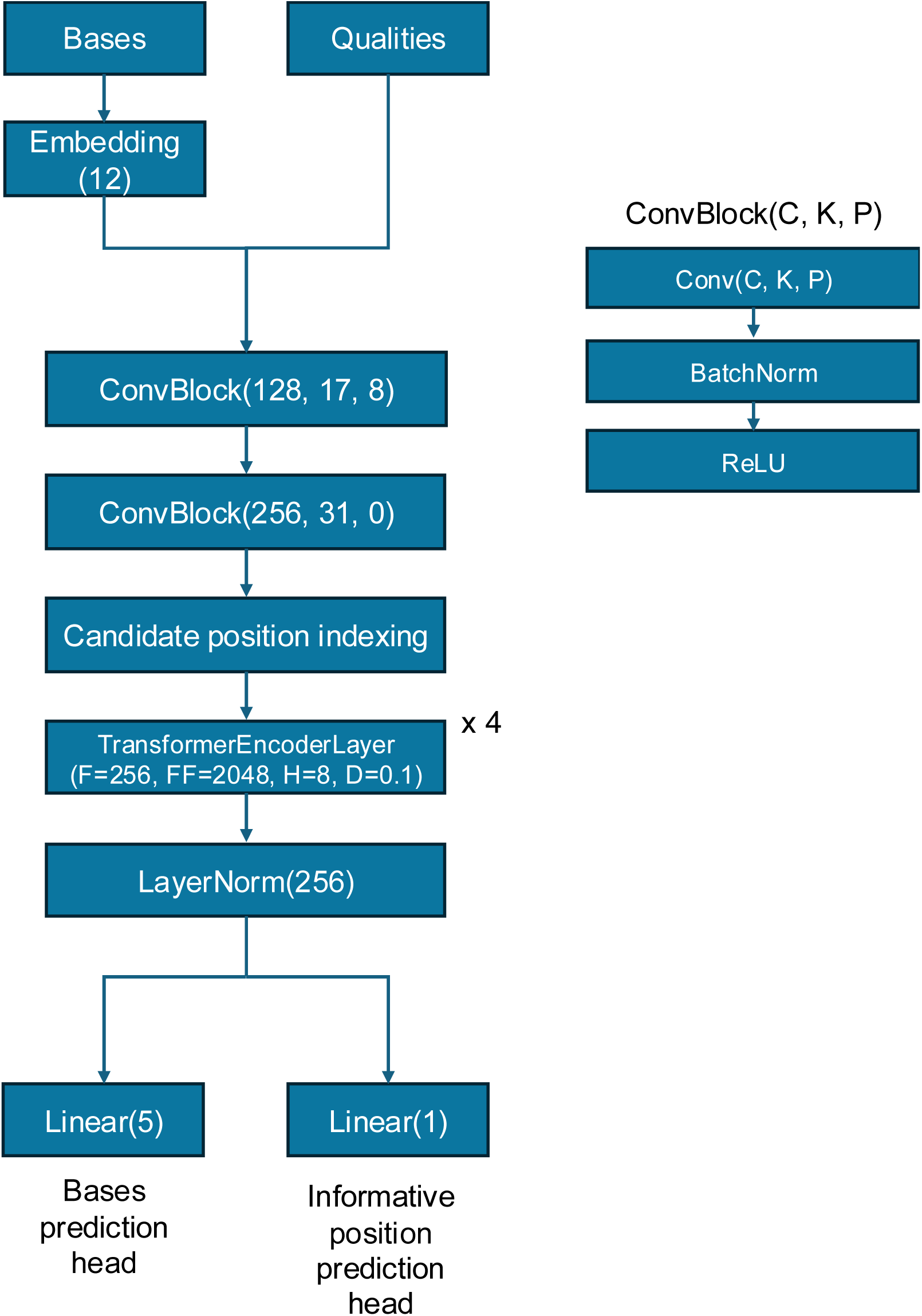
HERRO Model Architecture: The block diagram displays the complete architecture of the HERRO model, including layers and hyperparameters. The convolutional block, which consists of a convolutional layer, batch normalization, and a ReLU activation function, is fully defined on the right side of the figure.

## References

1. Nurk, S. et al. The complete sequence of a human genome. Science 376, 44–53 (2022).

2. Koren, S., et al. Gapless assembly of complete human and plant chromosomes using only nanopore sequencing. bioRxivorg (2024) doi:10.1101/2024.03.15.585294.

3. Sarashetti, P., Lipovac, J., Tomas, F., Šikic, M. & Liu, J. The hitchhiker’s guide to sequencing data types and volumes for population-scale pangenome construction. bioRxiv (2024) doi:10.1101/2024.03.14.585029.

4. Koren, S. et al. De novo assembly of haplotype-resolved genomes with trio binning. Nat. Biotechnol. (2018) doi:10.1038/nbt.4277.

5. Sanders, A. D., Falconer, E., Hills, M., Spierings, D. C. J. & Lansdorp, P. M. Single-cell template strand sequencing by Strand-seq enables the characterization of individual homologs. Nat. Protoc. 12, 1151–1176 (2017).

6. Koren, S., Walenz, B. P., Berlin, K., Miller, J. R. & Phillippy, A. M. Canu: scalable and accurate long-read assembly via adaptive k-mer weighting and repeat separation. bioRxiv, 071282. Genome Research 27, 722–736 (2017).

7. Guan, D. et al. Identifying and removing haplotypic duplication in primary genome assemblies. Bioinformatics 36, 2896–2898 (2020).

8. Chin, C.-S. et al. Phased diploid genome assembly with single-molecule real-time sequencing. Nat. Methods 13, 1050–1054 (2016).

9. Kolmogorov, M., Yuan, J., Lin, Y. & Pevzner, P. A. Assembly of long, error-prone reads using repeat graphs. Nat. Biotechnol. 37, 540–546 (2019).

10. Shafin, K. et al. Haplotype-aware variant calling with PEPPER-Margin-DeepVariant enables high accuracy in nanopore long-reads. Nat. Methods 18, 1322–1332 (2021).

11. Kolmogorov, M. et al. Scalable Nanopore sequencing of human genomes provides a comprehensive view of haplotype-resolved variation and methylation. Nat. Methods 20, 1483–1492 (2023).

12. Shafin, K. et al. Nanopore sequencing and the Shasta toolkit enable efficient de novo assembly of eleven human genomes. Nat. Biotechnol. 38, 1044–1053 (2020).

13. Nie, F. et al. De novo diploid genome assembly using long noisy reads. Nat. Commun. 15, 2964 (2024).

14. Cheng, H., Concepcion, G. T., Feng, X., Zhang, H. & Li, H. Haplotype-resolved de novo assembly using phased assembly graphs with hifiasm. Nat. Methods 18, 170–175 (2021).

15. Cheng, H. et al. Haplotype-resolved assembly of diploid genomes without parental data. Nat. Biotechnol. 40, 1332–1335 (2022).

16. Cheng, H., Asri, M., Lucas, J., Koren, S. & Li, H. Scalable telomere-to-telomere assembly for diploid and polyploid genomes with double graph. Nat. Methods (2024) doi:10.1038/s41592-024-02269-8.

17. Rautiainen, M. et al. Telomere-to-telomere assembly of diploid chromosomes with Verkko. Nat. Biotechnol. 41, 1474–1482 (2023).

18. Hu, J. et al. NextDenovo: an efficient error correction and accurate assembly tool for noisy long reads. Genome Biol. 25, 107 (2024).

19. Darian, J. C., Kundu, R., Rajaby, R. & Sung, W.-K. Constructing telomere-to-telomere diploid genome by polishing haploid nanopore-based assembly. Nat. Methods 21, 574–583 (2024).

20. Bankevich, A., Bzikadze, A. V., Kolmogorov, M., Antipov, D. & Pevzner, P. A. Multiplex de Bruijn graphs enable genome assembly from long, high-fidelity reads. Nat. Biotechnol. 40, 1075–1081 (2022).

21. Li, H. Minimap2: pairwise alignment for nucleotide sequences. Bioinformatics 34, 3094–3100 (2018).

22. Rhie, A. et al. Towards complete and error-free genome assemblies of all vertebrate species. Nature 592, 737–746 (2021).

23. Wang, B. et al. High-quality Arabidopsis thaliana genome assembly with nanopore and HiFi long reads. Genomics Proteomics Bioinformatics 20, 4–13 (2022).

24. Wenger, A. M. et al. Accurate circular consensus long-read sequencing improves variant detection and assembly of a human genome. Nat. Biotechnol. 37, 1155–1162 (2019).

25. Chernyavskaya, Y., Zhang, X., Liu, J. & Blackburn, J. Long-read sequencing of the zebrafish genome reorganizes genomic architecture. BMC Genomics 23, 116 (2022).

26. Kamath, S. S., Bindra, M., Pal, D. & Jain, C. Telomere-to-telomere assembly by preserving contained reads. bioRxiv (2023) doi:10.1101/2023.11.07.565066.

27. Jain, C. Coverage-preserving sparsification of overlap graphs for long-read assembly. Bioinformatics 39, (2023).

28. Li, H., Feng, X. & Chu, C. The design and construction of reference pangenome graphs with minigraph. Genome Biol. 21, 265 (2020).

29. Rhie, A., Walenz, B. P., Koren, S. & Phillippy, A. M. Merqury: reference-free quality, completeness, and phasing assessment for genome assemblies. Genome Biol. 21, 245 (2020).

30. Rhie, A. et al. The complete sequence of a human Y chromosome. Nature 621, 344–354 (2023).

31. Gurevich, A., Saveliev, V., Vyahhi, N. & Tesler, G. QUAST: quality assessment tool for genome assemblies. Bioinformatics 29, 1072–1075 (2013).

32. Yang, C. et al. The complete and fully-phased diploid genome of a male Han Chinese. Cell Res. 33, 745–761 (2023).

33. Pang, A. W. et al. Towards a comprehensive structural variation map of an individual human genome. Genome Biol. 11, R52 (2010).

34. Vollger, M. R. et al. Increased mutation and gene conversion within human segmental duplications. Nature 617, 325–334 (2023).

35. Logsdon, G. A. et al. Complex genetic variation in nearly complete human genomes. bioRxiv 2024.09.24.614721 (2024) doi:10.1101/2024.09.24.614721.

36. Vaser, R., Sović, I., Nagarajan, N. & Šikić, M. Fast and accurate de novo genome assembly from long uncorrected reads. Genome Res. 27, 737–746 (2017).

37. Nurk, S. et al. HiCanu: accurate assembly of segmental duplications, satellites, and allelic variants from high-fidelity long reads. Genome Res. 30, 1291–1305 (2020).

38. Šošic, M. & Šikic, M. Edlib: a C/C ++ library for fast, exact sequence alignment using edit distance. Bioinformatics 33, 1394–1395 (2017).

39. Groot Koerkamp, R. & Ivanov, P. Exact global alignment using A* with chaining seed heuristic and match pruning. Bioinformatics 40, (2024).

40. Kalikar, S., Jain, C., Vasimuddin, M. & Misra, S. Accelerating minimap2 for long-read sequencing applications on modern CPUs. Nat. Comput. Sci. 2, 78–83 (2022).

41. Sadasivan, H. et al. Accelerating Minimap2 for accurate long read alignment on GPUs. J. Biotechnol. Biomed. 6, 13–23 (2023).

42. Kolmogorov, M. et al. metaFlye: scalable long-read metagenome assembly using repeat graphs. Nat. Methods 17, 1103–1110 (2020).

43. Feng, X., Cheng, H., Portik, D. & Li, H. Metagenome assembly of high-fidelity long reads with hifiasm-meta. Nat. Methods 19, 671–674 (2022).

44. Bertrand, D. et al. Hybrid metagenomic assembly enables high-resolution analysis of resistance determinants and mobile elements in human microbiomes. Nat. Biotechnol. 37, 937–944 (2019).

45. Vaser, R. & Šikić, M. Time- and memory-efficient genome assembly with Raven. Nat. Comput. Sci. 1, 332–336 (2021).

46. Stanojević, D., Li, Z., Bakić, S., Foo, R. & Šikić, M. Rockfish: A transformer-based model for accurate 5-methylcytosine prediction from nanopore sequencing. Nat. Commun. 15, 5580 (2024).

47. Wick, R. R., Judd, L. M., Gorrie, C. L. & Holt, K. E. Completing bacterial genome assemblies with multiplex MinION sequencing. Microb. Genom. 3, e000132 (2017).

48. Shen, W., Sipos, B. & Zhao, L. SeqKit2: A Swiss Army Knife for Sequence and Alignment Processing. iMeta (2024).

49. Shen, W., Le, S., Li, Y. & Hu, F. SeqKit: A cross-platform and ultrafast toolkit for FASTA/Q file manipulation. PLoS One 11, e0163962 (2016).

50. Li, H. New strategies to improve minimap2 alignment accuracy. Bioinformatics 37, 4572–4574 (2021).

51. Vaswani, A., et al. Attention is all you need. arXiv [cs.CL] (2017).

52. Xiong, R., et al. On Layer Normalization in the Transformer Architecture. arXiv [cs.LG] (2020).

53. Manni, M., Berkeley, M. R., Seppey, M. & Zdobnov, E. M. BUSCO: Assessing genomic data quality and beyond. Curr. Protoc. 1, e323 (2021).

54. Manni, M., Berkeley, M. R., Seppey, M., Simão, F. A. & Zdobnov, E. M. BUSCO update: Novel and streamlined workflows along with broader and deeper phylogenetic coverage for scoring of eukaryotic, prokaryotic, and viral genomes. Mol. Biol. Evol. 38, 4647–4654 (2021).

55. Huang, N. & Li, H. compleasm: a faster and more accurate reimplementation of BUSCO. Bioinformatics 39, (2023).

56. Danecek, P. et al. Twelve years of SAMtools and BCFtools. Gigascience 10, (2021).

57. Bonfield, J. K. et al. HTSlib: C library for reading/writing high-throughput sequencing data. Gigascience 10, (2021).

58. Mason, J. M. & Biessmann, H. The unusual telomeres of Drosophila. Trends Genet. 11, 58–62 (1995).

