## Supplementary Information for "Telomere-to-Telomere Phased Genome Assembly Using HERRO-Corrected Simplex Nanopore Reads"

### Rebasecalling:

HG002 data was downloaded and used as it was - it was basecalled using Dorado v0.4.0 and model dna_r10.4.1_e8.2_400bps_sup@v4.1.0.
I002C data basecalled using Dorado v0.3.0 and model dna_r10.4.1_e8.2_400bps_sup@v4.1.0.

All rebasecalling was done using Dorado v0.5.3:
- For R9.4.1 data (CHM13), the model used was “dna_r9.4.1_e8_sup@v3.6”
- For R10.4.1 data (HG005, Drosophila melanogaster, HG02818, Danio rerio), the model used was dna_r10.4.1_e8.2_400bps_sup@v4.1.0.

1. Single-read FAST5 files were converted to multi-read FAST5 files for CHM13 using the single_to_multi_fast5 script from the ONT fast5 API software v4.1.3 (https://github.com/nanoporetech/ont_fast5_api):

```
single_to_multi_fast5 -i single_fast5_directory --recursive -t
$num_threads -s multi_fast5_directory
```

2. The multi-read FAST5 files were then converted to POD5 files for all rebasecalled samples using pod5 package v0.3.6 (https://github.com/nanoporetech/pod5-file-format):

```
pod5 convert fast5 -o converted.pod5 -t $num_threads -r input_dir
```

3. Basecalling:
   a. For all samples except CHM13:

```
dorado basecaller --emit-sam $model $pod5_folder > output.sam
```

   b. For CHM13:

```
dorado basecaller -r --emit-fastq $model $pod5_folder > output.fastq
```

For CHM13, FASTQ was extracted from SAM, and no other information from the SAM file was used.

### Correction using Herro:

The version used for this manuscript can be found at https://github.com/lbcb-sci/herro/tree/manuscript. The libtorch version used was https://download.pytorch.org/libtorch/cu118/libtorch-cxx11-abi-shared-with-deps-2.0.1%2Bcu118.zip . The Herro binary and preprocessing scripts used follow those in lbcb-sci/herro at commit c8cc4f87be6b5233db7ea80fb3206c4d726402d5 (https://github.com/lbcb-sci/herro/tree/c8cc4f87be6b5233db7ea80fb3206c4d726402d5) with two minor changes:

- INFER_CHANNEL_CAP_FACTOR in src/lib.rs was increased from 2 to 4.
- The length filtering step at the end of scripts/preprocess.sh and scripts/no_split.sh was removed, making the output from the previous step the final output.

- Steps 1. and 2. below were not done for reads basecalled using Dorado v0.5.0 and newer which integrates these steps into basecalling. They were done for the HG002 and I002C reads. The steps were performed on the HG002 Q28 reads before it was corrected. However, the untrimmed/split inputs were used whenever the corrected version was used (as UL input, or for the assembly of the uncorrected reads).

1. Adapter removal using Porechop (v0.2.4):

```
porechop --threads $num_threads -i input.fastq.gz --format fastq.gz -o
output.fastq.gz --adapter_threshold 95
```

2. Read splitting on adapters using duplex_tools (v0.3.3):

```
duplex_tools split_on_adapter --threads $num_threads
--allow_multiple_splits $input_dir $output_dir Native
```

3. Filtering short (<10 kbps) and low-quality (mean q-score < 10) using SeqKit (v2.5.1):

```
seqkit seq -m 10000 -Q 10 input.fastq.gz -o output.fastq.gz
```

4. All-vs-all alignments between reads were generated using the create_batch_alignments script (https://github.com/lbcb-sci/herro/blob/c8cc4f87be6b5233db7ea80fb3206c4d726402

d5/scripts/create_batched_alignments.sh). The minimap2 version used was 2.26-r1175. SeqKit v2.5.1 was used to obtain ids of the input reads:

```
seqkit seq -n -i input.fastq.gz > rids.txt
./create_batched_alignments.sh input.fastq.gz rids.txt $num_threads
$output_dir
```

**Note: The parameters for used for minimap2 in create_batched_alignments.sh was:**

```
minimap2 -K8g -cx ava-ont -k25 -w17 -e200 -r150 -m4000 -z200
-t${num_threads} --dual=yes $reads $reads
```

**and it was used for HG002, HG02818, HG005, CHM13, I002C and Danio rerio.**

**We used alternative parameters which we found to be more suitable for non-UL reads for A. thaliana, D. melanogaster, and an additional correction run for HG005:**

```
minimap2 -K8g -cx ava-ont -k21 -w14 -f 0.005 -e100 -r150 -z200 -m1500
-t${num_threads} --dual=yes $reads $reads
```

5. Run the HERRO binary:
    - The model for R10.4.1 can be downloaded from: http://complex.zesoi.fer.hr/data/downloads/model_v0.1.pt
    - The model for R9.4.1 can be downloaded from: http://complex.zesoi.fer.hr/data/downloads/model_R9_v0.1.pt

```
herro inference --read-alns $alns_dir -t $num_threads -d $gpus -m $model
-b 128 trimmed_and_filtered_reads.fastq.gz corrected_output.fasta
```

### Assembling:

**Subsampling of Reads for Assembly:**

The output reads from correction were downsampled to approximately 35× coverage to be used as a highly accurate/HiFi input substitute for Hifiasm/Verkko/LJA if the coverage was significantly higher than 35× (>38×). SeqKit v2.5.1 was used for downsampling.
Coverage was estimated by dividing the total number of base pairs by the estimated haploid genome size. The estimated haploid genome sizes used are:

- Human: 3.1G
- Danio rerio: 1.5G
- Arabidopsis thaliana: 135M
- Drosophila melanogaster: 160M

```
seqkit sample -p $proportion corrected.fasta -o downsampled.fasta
```

For some assembly runs input reads were filtered by length using SeqKit v2.5.1:

```
seqkit seq -m $min_length $input_fasta -o $output_fasta
```

**Hifiasm v0.19.8:**

Corrected reads were first chopped to a length of 30,000 bp, and chunks shorter than 10,000 bp were filtered away using SeqKit v2.5.1:

```
seqkit sliding -s 30000 -W 30000 -g corrected_reads.fasta >
chopped.fasta
seqkit seq -m 10000 chopped.fasta > processed.fasta
```

a. For haploid inputs, the following command is used:

```
hifiasm -t $num_threads -o output_prefix -l0 -ul $ultra_long_input
--ul-cut 50000 processed.fasta
```

b. Creation of yak files from short reads for trio-binning mode of Hifiasm for diploid inputs: yak 0.1-r69-dirty was used for all samples except for I002C which used 0.1-r66-dirty:

```
yak count -b37 -t $num_threads -o output.yak <(zcat sr*.fastq.gz) <(zcat
sr*.fastq.gz)
```

c. For diploid inputs (with parental data), the following command is used,

```
hifiasm -t $num_threads -o output_prefix --ul $ultra_long_input --ul-cut
50000 --dual-scaf -1 parent1.yak -2 parent2.yak processed.fasta
```

**Verkko:**

Verkko (v2.0) was used for most of the inputs. A few of the inputs crashed with that version. For these inputs, Verkko's master branch commit b050ef42e33ce0ac5bd806746172ae15c4c2dcae (https://github.com/marbl/verkko/tree/b050ef42e33ce0ac5bd806746172ae15c4c2dcae) along with MBG Branch master commit 10782bdeb38e6cd33918479e3f13413173aeb284 (https://github.com/maickrau/MBG/tree/10782bdeb38e6cd33918479e3f13413173aeb284) was used instead. The specific runs that crashed and used the alternative version instead are marked with "Verkko*" in the supplementary tables with assembly results. For more explanations, please refer to "Note on headings" in the explanation for supplementary tables. The $ram for all runs was 400.

a. For haploid inputs:

```
verkko -d output_directory --hifi highly_accurate_input.fasta --nano
ultra_long_input.fastq --local-memory $ram --local-cpus $num_cpus
--cleanup --haploid
```

b. To create hampers for trio mode of Verkko for diploid inputs:

Run the following for the assembly sample and the parents' short reads, this step was done with Meryl 1.4.1 for all samples except for I002C which used Meryl 1.4

```
meryl count compress k=30 threads=$num_threads memory=$ram *.fastq.gz
```

```
output prefix_compressed.k30.meryl
```

Create hapmers for verkko assembly, Merqury commit 4b4846a78e8eaa06e42e06ec9f40c7f329e664c4 (https://github.com/marbl/merqury/tree/4b4846a78e8eaa06e42e06ec9f40c7f329e664c4) with Meryl 1.4.1 was used for all samples for this step

```
$MERQURY_DIRECTORY/trio/hapmers.sh maternal_compressed.k30.meryl
paternal_compressed.k30.meryl child_compressed.k30.meryl
```

c. For diploid inputs (with parental data):

```
verkko -d output_directory --hifi highly_accurate_input.fasta --nano
$ultra_long_input --local-memory $ram --local-cpus $num_cpus --cleanup
--hap-kmers maternal_compress.k30.hapmer.meryl
paternal_compress.k30.hapmer.meryl trio
```

d. For assembling of HG002 corrected reads without parental data (for the X and Y contigs), the alternative version was used since v2.0 crashed for the run:

```
verkko -d output_directory --hifi highly_accurate_input.fasta --nano
ultra_long_input.fastq --local-memory $ram --local-cpus $num_cpus
--cleanup
```

**LJA (v0.2) - haploid/homozygous samples only:**

```
lja -o output_directory --reads input.fasta -t $num_threads
```

### Evaluation:

References used for evaluation:

- HG002 - hg002 v1.0.1 (https://github.com/marbl/HG002)
- HG02818 - chm13 v2.0 (https://github.com/marbl/CHM13)
- HG005 - chm13 v2.0
- CHM13 - chm13 v2.0
- I002C - I002C v0.4 (https://github.com/LHG-GG/I002C or https://github.com/lbcb-sci/I002C)
- Danio rerio - ASM3317019v2 (https://www.ncbi.nlm.nih.gov/datasets/genome/GCA_033170195.2/)
- Arabidopsis thaliana - Col-XJTU (https://ngdc.cncb.ac.cn/gwh/Assembly/21820/show)
- Drosophila melanogaster - ASM1890436v1 (https://www.ncbi.nlm.nih.gov/datasets/genome/GCA_018904365.1/)

The only exception is asmgene assessment where chm13 v2.0 is used for all human datasets.

**Assembly assessment using minigraph and paftools:** minigraph 0.20-r574-dirty, SAMtools 1.19.2 (Using htslib 1.19.1), paftools 2.26-r1175 were used.

```
minigraph -xasm --show-unmap=yes -t $num_threads reference.fasta \
    assembly.fasta > alignment.paf
samtools faidx reference.fasta
paftools.js asmstat reference.fasta.fai alignment.paf > output.txt
```

**Assembly assessment using yak**

a. Creation of yak files from short reads: yak 0.1-r69-dirty was used for all samples except for I002C which used 0.1-r66-dirty:

```
yak count -b37 -t $num_threads -o output.yak <(zcat sr*.fastq.gz) <(zcat
sr*.fastq.gz)
```

b. Calculating yak (0.1-r69-dirty) QV:

```
yak qv -t $num_threads child.yak assembly.fasta > output.txt
```

c. Calculating hamming and switch errors using yak (0.1-r69-dirty) trioeval:

```
yak trioeval -e -t $num_threads parent1.yak parent2.yak \
haplotype1_assembly.fasta > haplotype1.trioeval.txt
yak trioeval -e -t $num_threads parent1.yak parent2.yak \
haplotype2_assembly.fasta  > haplotype2.trioeval.txt
```

**Assembly assessment using Merqury:**
Commit 4b4846a78e8eaa06e42e06ec9f40c7f329e664c4 (https://github.com/marbl/merqury/tree/4b4846a78e8eaa06e42e06ec9f40c7f329e664c4) with with Meryl 1.4.1:

a. Build k-mer dbs with meryl 1.4.1

```
meryl count k=$k threads=$num_threads memory=$ram *.fastq.gz output
$output_prefix.meryl
```

b. Build hap-mers

```
$MERQURY_DIR/trio/hapmers.sh maternal.meryl paternal.meryl child.meryl
```

c. Evaluation for the case when it is a diploid sample and there's parental data:

```
OMP_NUM_THREADS=$num_threads $MERQURY_DIR/merqury.sh child.meryl
maternal.hapmer.meryl paternal.hapmer.meryl hap1.fasta hap2.fasta
$output_prefix
```

d. Hamming error rates were obtained by, taking the last column of the output (MajorHapError(%)):

```
$MERQURY_DIR/trio/hamming_error.sh $output_prefix.hapmers.count
hap1.fasta hap2.fasta
```

e. Evaluation for the case when it is a haploid/homozygous sample and there's no parental data:

```
OMP_NUM_THREADS=$num_threads $MERQURY_DIR/merqury.sh child.meryl
assembly.fasta $output_prefix
```

We use different k values for different organisms (obtained using $MERQURY_DIR/best_k.sh):

- Human - 21
- Danio rerio - 20
- Drosophila melanogaster - 19
- Arabidopsis thaliana - 18

**T2T assessment:**

a. The script from https://github.com/prasad693/Tel_Sequences (main branch commit 1a20c51620206fac666155263786cbc566366fe3) was used to find T2T contigs/scaffolds:

```
T2T_chromosomes.sh -a assembly.fasta -r reference.fasta -o output_prefix
-m $telomere_motif -t $num_threads
```

b. seqtk 1.4-r130-dirty was used to identify sequences with gaps and determine which of the T2T sequences were scaffolds.

```
seqtk gap assembly.fasta > gaps.tsv
cut -f1 gaps.tsv > ids_with_gaps.txt
grep -w -f ids_with_gaps.txt output_prefix_alignment_T2T.txt >
scaffolds_ids.txt
grep -w -v -f ids_with_gaps.txt output_prefix_alignment_T2T.txt >
contigs_ids.txt
```

The above was executed for each haplotype and its reference for diploid samples.

The telomere_motif used for each sample: humans - TTAGGG, Danio rerio - TTAGGG, Arabidopsis thaliana - TTTAGGG. Finding T2T was not done for Drosophila melanogaster.

Note: For LJA assemblies, assembly contigs were renamed to avoid an issue with contigs with different name lengths in the T2T assessment script. SeqKit v2.5.1 was used:

```
seqkit replace -p .+ -r "ctg{nr}" --nr-width 8 \
    assembly.fasta > assembly.renamed.fasta
```

**Asmgene for evaluating gene completeness and duplication**: minimap2 2.26-r1175 and paftools 2.26-r1175 were used.

```
minimap2 -cxsplice:hq -t $num_threads reference.fasta \
    cdna.fasta > ref.paf

minimap2 -cxsplace:hq -t $num_threads assembly.fasta \
    cdna.fasta > assembly.paf

paftools.js asmgene -a -i.97 ref.paf assembly.paf > asmgene_output.txt
```

The above was executed for each haplotype and its reference for diploid samples.

cDNA data used:

- Human - https://ftp.ensembl.org/pub/release-111/fasta/homo_sapiens/cdna/Homo_sapiens.GRCh38.cdna.all.fa.gz
- Drosophila melanogaster - https://ftp.ensembl.org/pub/release-111/fasta/drosophila_melanogaster/cdna/Drosophila_melanogaster.BDGP6.46.cdna.all.fa.gz
- Arabidopsis thaliana - https://ftp.ensemblgenomes.ebi.ac.uk/pub/plants/release-58/fasta/arabidopsis_thaliana/cdna/Arabidopsis_thaliana.TAIR10.cdna.all.fa.gz
- Danio rerio - https://ftp.ensembl.org/pub/release-111/fasta/danio_rerio/cdna/Danio_rerio.GRCz11.cdna.all.fa.gz

**Busco and compleasm evaluation**
BUSCO 5.6.1 and compleasm 0.2.5 were used.

```
busco -i assembly.fasta -m genome -o output_label -l $lineage \
    -c num_threads --download_path $busco_library

compleasm run -t $num_threads -l $lineage -L $compleasm_library \
    -a assembly.fasta -o output_prefix
```

Lineages (all dated 2024-01-08):

- Human - primates_odb10
- Drosophila melanogaster - diptera_odb10
- Danio rerio - actinopterygii_odb10
- Arabidopsis thaliana - brassicales_odb10

**Assessment of assembly with Quast:**

1. evaluation against reference with Quast 5.2.0, with minimap2 2.24-r1122,

```
quast -t $num_threads -r reference.fasta -o $output_prefix
assembly.fasta
```

**Other assembly assessment commands:**

2. calN50 (https://github.com/lh3/calN50) version r4 (k8 - V8: 3.16.4, K8: 0.2.5-r80) for calculating N50 and L90,

```
calN50.js assembly.fasta > output.txt
```

The above was executed for each haplotype for diploid samples.

3. Counts of Ns in scaffolded assemblies (SeqKit v2.5.1):

```
seqkit stats -a -G 'N' assembly.fasta > stats.txt
```

The above was executed for each haplotype for diploid samples.

**Evaluation of reads using bamConcordance:**

a. Reads were mapped with minimap2 2.26-r1175, SAMtools 1.19.2 (Using htslib 1.19.1) was used for converting to bam, sorting and indexing.

```
minimap2 -a -Q --eqx --secondary=no -I9G -K8G -t $num_threads \
    reference.fasta reads.fasta | samtools view -Sb \
    -@$num_threads > mappings.bam

samtools sort -@$num_threads mappings.bam -o mappings.sorted.bam
samtools index -@$num_threads mappings.sorted.bam
```

b. BAM file is used as input to bamConcordance (https://github.com/PacificBiosciences/hg002-ccs/blob/master/concordance/bamConcordance), pysam 0.22.0 (https://github.com/pysam-developers/pysam) was used:

```
bamConcordance reference.fasta mappings.bam out.csv
```

**Reads assessment using yak**

a. Creation of yak files from short reads: yak 0.1-r69-dirty was used for all samples except for I002C which used 0.1-r66-dirty:

```
yak count -b37 -t $num_threads -o output.yak <(zcat sr*.fastq.gz) <(zcat sr*.fastq.gz)
```

b. Calculating yak (0.1-r69-dirty) QV:

```
yak qv -t $num_threads child.yak reads.fastx > output.txt
```

The adjusted_quality_value in the output was used.

**Corrected reads' error rates estimation against references**

a. Corrected reads were subsampled to 20x using Seqkit V2.5.1 to reduce mapping time required.

```
seqkit sample -p $proportion corrected.fasta -o downsampled.fasta
```

b. Reads were mapped to the reference fasta. Reads were mapped with minimap2 2.26-r1175, SAMtools 1.19.2 (Using htslib 1.19.1) was used for converting to bam, sorting and indexing:

```
minimap2 -a --secondary=no -I9G -K8G -t$num_threads reference.fasta
corrected_reads.20x.fasta | $samtools view -Sb -@$num_threads >
mapping.bam
```

c. https://github.com/lbcb-sci/herro/blob/manuscript/scripts/error_rates.py was used to estimate error rates, pysam 0.22.0 (https://github.com/pysam-developers/pysam) was used:

```
error_rates.py -b mapping.bam
```

### CHM13 data partitions used

Some partitions from https://s3-us-west-2.amazonaws.com/human-pangenomics/index.html?prefix=T2T/CHM13/nanopore/fast5/ were skipped due to reasons such as having too many fast5 files. The partitions used are: 001-097, 101, 103-144, 149-152, 158-174, 177-178, 180-197, 202-204, 207-208, 213-215, 224 and 225 (partial).

### Additional notes on Arabidopsis thaliana data

Extraction protocol:
https://community.nanoporetech.com/extraction_methods/arabidopsis-leaf-dna
Library preparation protocol:
https://community.nanoporetech.com/docs/prepare/library_prep_protocols/genomic-dna-by-ligation-sqk-lsk114/v/gde_9161_v114_revu_29jun2022
